# Chromosome-Scale Assemblies of Flowering Dogwood Cultivars Enable Identification of Candidate Genes Regulating Anthocyanin Biosynthesis in Leaves and Bracts

**DOI:** 10.1101/2025.07.31.667920

**Authors:** Trinity P. Hamm, Sarah L. Boggess, Marcin Nowicki, Denita Hadziabdic, DeWayne Shoemaker, Robert N. Trigiano, William E. Klingeman, Thomas J. Molnar, Jim Leebens-Mack, Alex Harkess, Sheron A. Simpson, Ramey C. Youngblood, Amanda M. Hulse-Kemp, Brian E. Scheffler, Margaret E. Staton

## Abstract

- The North American-native ornamental tree, flowering dogwood (*Cornus florida* L.), has a showy bract display that can range in color from white to pink to deep red. Although many trees have white bracts, there is consumer demand for novel pigmentation in the bracts combined with other traits of interest. Because the genetic basis of all traits in flowering dogwood is unknown, combining them using traditional breeding efforts is time, labor, and space-intensive.
- We developed foundational genomic resources to establish marker-assisted selection within flowering dogwood breeding. We generated diploid, chromosome-scale, annotated genome assemblies for one pink-bracted and red-leafed tree and one white-bracted and green-leafed tree. Additionally, a phenotyping protocol for bract color and presence/absence diagnostic SNPs for bract and leaf color were established.
- We leveraged these resources to evaluate linkage associations and differential gene expression related to anthocyanin biosynthesis to identify candidate genes regulating bract and leaf pigmentation. Within a 14Mb locus we identified 14 anthocyanin-related candidate genes. Two genes, with MYB (*g19533*) and RING finger (*g19556)* binding domains, had both differential gene expression and variants with the expected segregation pattern.
- These resources will be valuable in combining pink-red bracts with other traits to advance flowering dogwood breeding.

## Introduction

Flowering dogwoods (*Cornus florida* L.) are small deciduous trees native to eastern North America, with the species native range stretching from Canada to Mexico (Eyde, 1988). Widely favored as ornamental plants in the United States (U.S.), they offer true four-season interest—displaying striking bracts in spring, vibrant red foliage in autumn, vivid red drupes through winter, and elegant structural form throughout the year (Cappiello & Shadow, 2005). In 2019, dogwoods accounted for over $31 million (USD) in wholesale and retail sales from 1.2 million trees, making them the third most valuable deciduous flowering tree in the U.S. market (USDA-NASS, 2020).

While dogwoods offer year-round beauty, the trees are most celebrated for their striking spring display. The captivating spring color of flowering dogwood originates from the floral bracts that subtend the small, true flowers. Bract color can range from white to pink to deep red, varying by cultivar and environment (Orton, 1982; Cappiello & Shadow, 2005). The bracts can also display impressive patterning both within and around the veins. The pink-red pigmentation can be induced or deepened on bracts when infected by *Elsinoe corni* (a fungus that causes spot anthracnose) on both white and pink bracts.

The new leaves of flowering dogwood cultivars range in color from green to red, typically matching the color and intensity of the bracts and adding to the ornamental interest (Orton, 1982). Two popular cultivars are ‘Cherokee Brave’, which has pink bracts and red leaves, and ‘Appalachian Spring’, which features white bracts and green leaves. Both of these cultivars are susceptible to powdery mildew, which is caused by the fungus *Erysiphe pulchra*. Currently, flowering dogwood breeding efforts are primarily focused on combining pink-red bracts and leaves with resistance to powdery mildew (Molnar & Capik, 2013; Molnar, 2022).

Pink-red pigmentation in plants, regardless of tissue type, is typically caused by anthocyanins (see reviews: Tanaka et al., 2008; Narbona et al., 2021). This is also the case in the vegetative and floral tissues of flowering dogwoods (Wadl *et al*., 2011). Anthocyanins are a class of flavonoids and are suggested to protect against UV radiation, herbivory, and pathogens as well as enhance tolerance to drought, cold, high salinity, and nutrient deficiency (LaFountain & Yuan, 2021; Günther *et al*., 2022; Li & Ahammed, 2023). The production of anthocyanins occurs in three main stages: the general phenylpropanoid pathway, the flavonoid pathway, and finally, the anthocyanin-specific pathway. Within the anthocyanin-specific pathway, three branches lead to the production of cyanidin, pelargonidin, and delphinidin derivatives. The combinations of these compounds are responsible for the wide range of reds, blues, and purples hues in flowering plants (Naing & Kim, 2018; Narbona *et al*., 2021; Bouillon *et al*., 2024).

The anthocyanin regulatory network is composed of structural genes that encode enzymes catalyzing pigment formation and regulatory elements that modulate the structural enzyme expression (Kodama *et al*., 2018). The main transcription factors that regulate the expression of the structural enzymes are classified into three categories based on their DNA binding domains: R2R3-MYB, basic helix-loop-helix (bHLH), and conserved WD40 repeats (WDR). Collectively, this MYB-bHLH-WDR regulatory complex (MBW complex) activates multiple genes in the anthocyanin production pathway (LaFountain & Yuan, 2021). This MBW activating complex is highly conserved in flowering plants (An *et al*., 2020; LaFountain & Yuan, 2021).

Regulation of the anthocyanin biosynthetic pathway can occur within any of the three main stages, and R2R3-MYBs are some of the most important and versatile transcription factors involved. R2R3-MYBs can promote or repress the expression of multiple enzymes throughout any of the three stages of the pathway (Schwinn *et al*., 2006; Albert *et al*., 2011). Within the first two stages, R2R3-MYBs can play a role in the competition of other secondary metabolite pathways by regulating the expression of structural genes that control the flux of precursor compounds throughout the various secondary metabolite pathways (Albert *et al*., 2011; Yuan *et al*., 2016). For example, in monkey flowers (*Mimulus* spp.), an R2R3-MYB (*LAR1*) has been shown to positively regulate *FLAVONOL SYNTHASE* (*FLS*), which is part of the flavonoid biosynthesis pathway and diverts precursor anthocyanin compounds to flavonol biosynthesis, causing areas of the flowers to be white (Yuan *et al*., 2016). Additionally, the effectiveness of all of these MYBs can be impacted by proteins like RING Zinc Fingers, which can either stabilize them (Yang *et al*., 2024) or mark them for degradation with ubiquitination (An *et al*., 2017).

The regulation of these pathways occurs on a spatial level, as the pigmentation accumulation can be present, absent, or occur in a gradient across specific parts of a tissue (Yuan *et al*., 2016; Kodama *et al*., 2018). R2R3-MYBs have been found to impact the pigmentation patterning in snapdragons (*Antirrhinum*; Schwinn et al., 2006; Li et al., 2016), petunia (*Petunia*; Albert et al., 2011), and monkey flower (*Mimulus*; Yuan et al., 2016). In the petunia study, separate R2R3-MYB transcription factors were identified as regulators of vein-associated pigmentation (*DPL*) and bud-blush (*PHZ*) that results in a tie-dye appearance on the petals when exposed to light (Albert *et al*., 2011).

The anthocyanin biosynthetic pathway also relies on and is impacted by external stimuli, including plant hormones such as auxin. The homeostasis of auxin is controlled by the vacuolar auxin transport mediator, Walls Are Thin 1 (*WAT1*) (Ranocha *et al*., 2013), and knocking out the gene causes anthocyanin accumulation in tomatoes (*Solanum lycopersicum*) (Koseoglou *et al*., 2023). While the exact mechanism for *WAT1* impacting anthocyanin accumulation is unknown, there is greater understanding of how other auxin-related mechanisms regulate anthocyanin accumulation. For example, in apple (*Malus domestica*) auxin regulates anthocyanin biosynthesis with an Aux/IAA repressor (*MdIAA121*) and an Auxin Response Factor (*MdARF13*) (Wang *et al*., 2018).

Research on anthocyanin production in *Cornus* species has been limited. Previous anthocyanin profiling studies have only included fruits and did not investigate the leaves or bracts (Du *et al*., 1974; Vareed *et al*., 2006). The red color in the drupes of flowering dogwood is mostly caused by the pigments Cyanidin 3-*O*-galactoside and Cyanidin-*O*-glucoside, with low levels of Delphinidin 3-*O*-glucoside (Vareed *et al*., 2006). The sole genetic analysis of color in flowering dogwoods identified four putative quantitative trait loci (QTL) associated with leaf color, located on three linkage groups (Wadl *et al*., 2011). However, the QTL are based on sparsely dispersed simple sequence repeat (SSR) markers and a small population size (n=94). Additionally, the identified QTL explained low phenotypic variance (1-17%) and exhibited instability over the five years of phenotyping. To date, no QTL study has investigated bract color, although most flowering dogwood trees with pink-red bracts also have red leaves, and therefore, one hypothesis is that a single QTL is associated with both traits. Additionally, there was previously no chromosome-scale well-annotated reference genome available for flowering dogwood that could be used to give greater context to these results. Therefore, the limited understanding of the genetic basis of anthocyanin biosynthesis in dogwoods hinders its practical application in breeding efforts.

The goal of this study was to understand the regulation of anthocyanin biosynthesis in flowering dogwood, specifically control of the varying bract and leaf color in *C. florida* ‘Cherokee Brave’ and ‘Appalachian Spring’. To accomplish this, we: 1) *de novo* assembled and annotated genomes for the two flowering dogwood cultivar parents; 2) genotyped and phenotyped a “pseudo-F2” population (n=125); 3) identified one locus associated with bract and leaf color; and 4) identified differentially expressed genes in ‘Cherokee Brave’ and ‘Appalachian Spring’ in a time series RNAseq experiment. As a result, we identified several candidate genes within the 14Mb QTL, some with differential gene expression. We identified diagnostic single nucleotide polymorphism (SNPs) that can predict the presence/absence of bract and leaf color in our population. The genetic and genomic resources developed in this study will guide future research and breeding initiatives aimed at combining deeper pink-red bracts with pathogen resistance traits (e.g., *E. pulchra* resistance) for flowering dogwood.

## Materials and Methods

### Plant Materials and Sequencing–Genome Assembly

Flowering dogwood cultivars ‘Cherokee Brave’ and ‘Appalachian Spring’ were selected for sequencing and subsequent genome assembly. ‘Cherokee Brave’ has pink bracts and red leaves, whereas ‘Appalachian Spring’ has white bracts and green leaves. ‘Cherokee Brave’ flower buds were collected from a residential tree in Knoxville, TN, U.S., with permission from the landowner. Young leaves from ‘Appalachian Spring’ were collected on the campus of the University of Georgia. Following high molecular weight (HMW) DNA extraction and library preparation, the libraries were sequenced using HiFi technology on the PacBio Sequel II for ‘Appalachian Spring’ and Sequel IIe for ‘Cherokee Brave’ (Pacific Biosciences, Menlo Park, CA, U.S.). Additional information on material and methods can be found in Supporting Information Methods S1. The genome size of ‘Cherokee Brave’ was confirmed with flow cytometry. Additionally, young leaves were collected from ‘Appalachian Spring’ and ‘Cherokee Brave’ trees on the University of Tennessee, Knoxville campus for Illumina Hi-C sequencing at Phase Genomics (Seattle, WA, U.S.).

### Plant Materials and Sequencing–Genotyping

A “pseudo-F2” population was previously generated from ‘Cherokee Brave’ and ‘Appalachian Spring’ (Wang *et al*., 2009). Due to the self-incompatibility of flowering dogwood and a second generation being necessary to recover the segregating pink bract phenotype, two ‘Cherokee Brave’ × ‘Appalachian Spring’ F1s (97-7 and 97-6) were crossed to generate the “pseudo-F2” population (Wang *et al*., 2009). The resulting population segregates for bract and leaf color and remains at the University of Tennessee Arboretum (Oak Ridge, TN, U.S.). To genotype the population, young leaf material was collected from all 125 individuals in the population and DNA was extracted. Once quality was ensured, samples were sent for genotype-by-sequencing (GBS) at The University of Wisconsin-Madison Biotechnology Center (Madison, WI, U.S.) with *PstI*/*MspI* restriction enzymes.

### Plant Materials and Sequencing–RNAseq

To identify differentially expressed genes between the white and pink bracts as well as the green and red leaves, developing leaf and bract tissues were collected and immediately flash frozen in liquid nitrogen. Three replicates of tissues were collected from one ‘Cherokee Brave’ (located in a residential area in Knoxville, TN, U.S., collected with the permission of the landowner) and two ‘Appalachian Spring’ (located on the University of Tennessee, Knoxville campus, U.S.) trees over a time course (Figure 1a).

**Figure 1.**
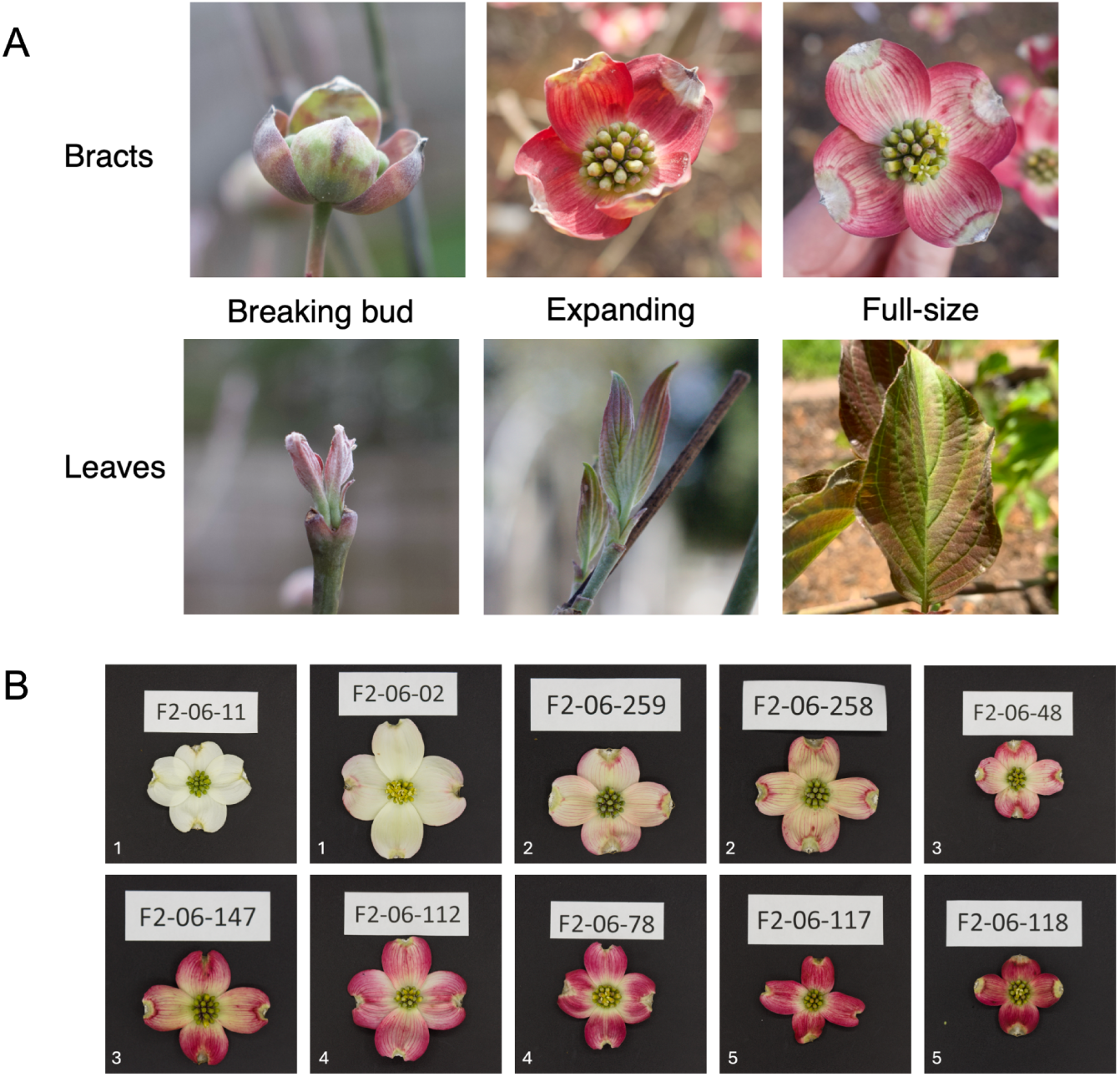
A) Developmental timepoints collected for leaves and bracts of *Cornus florida* ‘Cherokee Brave’ and ‘Appalachian Spring’ for the timecourse RNAseq experiment. Pictured is ‘Cherokee Brave’. B) Representative bract variation within the ‘Cherokee Brave’ × ‘Appalachian Spring’ “pseudo-F2” flowering dogwood population. White numbers in the lower left corner represent phenotypic class assigned (1 = white bract/green leaf, 2 = less than half pink bract/red leaf, 3 = half pink bract/red leaf, 4 = more than half pink bract/red leaf, 5 = almost completely pink bract/red leaf). Tree ID is on the white label within the image.

Thirty-six tissue samples were collected at the following three developmental timepoints: 1) breaking bud, 2) expanding tissue, and 3) fully expanded tissue. RNA was extracted from the tissues and sequenced.

### Genome Assembly and Scaffolding

Chromosome-scale, partially haplotype-resolved genome assemblies were completed for both ‘Cherokee Brave’ and ‘Appalachian Spring’ using default parameters of *hifiasm* 0.16.1-r375 (Cheng *et al*., 2021).

Quality was assessed with *BBTools* (Bushnell, 2022) and *BUSCO* 5.3.2 using the embryophyta_odb10 dataset run (Manni *et al*., 2021). Both the HiFi and Hi-C data were used as input to build this initial contig-level assembly. From there, Hi-C contacts were identified in the preliminary hifiasm assembly with *juicer* 1.6 (Durand *et al*., 2016b) and scaffolded into 11 chromosomes using *3D-DNA* v.201008 (Dudchenko *et al*., 2017). The genome was numbered and oriented against an existing linkage map for flowering dogwood (Pfarr Moreau *et al*., 2022) using NCBI *blast* 2.11.0 (Camacho *et al*., 2009) and *RIdeogram* 0.2.2 (Hao *et al*., 2020). Additionally, the percentage of markers aligning to the corresponding chromosome/linkager group were calculated. To subjectively assess the quality of the genomes, haplotypes 1 and 2 for each cultivar were compared to one another using *SyRI* 1.7.0 (Goel *et al*., 2019), and visualized with *plotsr* 1.1.1 (Goel & Schneeberger, 2022). The scaffolded assemblies were edited with *Juicebox* 1.7.0 (Durand *et al*., 2016a). Chloroplast and mitochondrial scaffolds were removed from the assembly based on alignment to the flowering dogwood chloroplast genome (NCBI: NC_044820.1) and rhododendron mitochondrial genome (NCBI: OQ450179.1). Additional contamination screens were completed using *FCS-gx* 0.4.0 and *FCS-adaptor* 0.5.0 (Astashyn *et al*., 2024).

### Genome Annotation

Repeats were identified with *RepeatModeler* 2.0.3 (Flynn *et al*., 2020) and masked using *RepeatMasker* 4.1.3-p1 (Smit *et al*., 2013). Genome annotation was completed with *BRAKER3* 3.0.3 (docker container v.1.0.3) (Gabriel *et al*., 2023) with both RNAseq data from the respective cultivars and closely related protein databases used as extrinsic evidence. *BUSCO*, *EnTAP* 1.0.0 (Hart *et al*., 2020), and *gFACs* (Caballero & Wegrzyn, 2019) were used to assess genome annotation.

### Population Genotyping

To genotype the 125 individuals in the “pseudo-F2” population, the raw GBS reads were demultiplexed with *sabre* 1.0 (Joshi, 2011), trimmed with *Fastp* 0.23.4 (Chen, 2023), and aligned to the ‘Cherokee Brave’ Haplotype 2 genome with *bwa* 0.7.17 (Li, 2013). Joint genotyping was completed with *GATK* 4.2.2.0 (Poplin *et al*., 2017), and filtered with *bcftools* 1.6 (Danecek *et al*., 2021) using a parameter sweep. One individual was removed due to failed sequencing (F2-06-65). The final filtering thresholds used were minimum read depth of 10, maximum missing rate of 20%, and minor allele frequency of 0.05. A kinship matrix was built with *plink2* 2.00 (Chang *et al*., 2015) and visualized with a heatmap in R with *ggplot* 3.5.1. Eight samples that had an average kinship value of <0 were removed from the dataset (F2-06-01, F2-06-120, F2-06-220A, F2-06-128, F2-06-63, F2-06-82, F2-06-60, and F2-06-219). This filtered VCF file was used for linkage map construction.

### Linkage Map Construction

Linkage map construction was completed with *OneMap* 3.0.0 (Margarido *et al*., 2007) using both the recombination and reference genome information of the individuals left after SNP and relatedness filtering (n=116). The tutorial vignette for outcrossing populations was followed, using ‘97-6’ as parent 1 and ‘97-7’ as parent 2.

### Population Phenotyping

The bracts of the “pseudo-F2” population were phenotyped between March 31 and April 6, 2023, at the UT Arboretum (Oak Ridge, TN, U.S.). Trees (n=125) were surveyed every day during this period, however, only 117 bloomed that year. For the bracts, multiple phenotyping methods were used. The inflorescences were photographed in a light box with a DSLR camera for subsequent image analysis. For each inflorescence collected, three images were captured (Figure S1): 1) full inflorescence, 2) four bracts pulled off of true flowers, and 3) one bract. The one bract in image 3 also had two colorimeter readings (in L*a*b* colorscale) taken with the CR-10 Plus Colorimeter (Konica Minolta, Chiyoda, Tokyo, Japan).

Categorical data was also collected, for both leaves (assessed on April 20, 2023) and bracts, using five classes (1 = white bract/green leaf, 2 = less than half pink bract/red leaf, 3 = half pink bract/red leaf, 4 = more than half pink bract/red leaf, 5 = almost completely pink bract/red leaf; Figure 1). The white balance of the images was corrected using RawTherapee (https://rawtherapee.com/), and image files were renamed to the tree ID using a custom python script (available https://github.com/trinityhamm/Flowering-Dogwood-Color). Images were then analyzed using a Fiji macro (Strock, 2021) to segment images to just the inflorescences or bracts and extract average RGB and L*a*b values.

### Anthocyanin Profiling

Qualitative anthocyanin profiling was performed on the bracts and leaves of ‘Cherokee Brave’ by Creative Proteomics (Shirley, NY, U.S.).

### QTL Mapping

All 23 phenotyping metrics (RGB and L*a*b* from full inflorescence, four bracts, and one bract images, L*a*b* from colorimeter, and five-class classification for the leaves and bracts) were used for composite interval mapping (CIM) with *fullsibQTL* 0.0.9012 (Gazaffi *et al*., 2020). A total of 116 individuals had leaf and genotypic data, and 108 individuals had bract, leaf, and genotypic data. The threshold for QTL detection was then determined for select methods with 1,000 permutations.

### Differential Gene Expression

Differentially expressed genes were identified between the white-bracted/green-leafed ‘Appalachian Spring’ and the pink-bracted/red-leafed ‘Cherokee Brave’. The reads were quality assessed with *fastqc* 0.12.1 (Andrews, 2010) and mapped to the ‘Cherokee Brave’ Hap 2 annotated genome with *STAR* 2.7.11b (Dobin *et al*., 2013). Transcripts were counted with *HTseq* 2.0.4 (Anders *et al*., 2015), and differentially expressed genes were identified with *DESeq2* 1.46.0 (Love *et al*., 2014). The likelihood ratio test (LRT) was used with a reduced model. Differentially expressed genes were identified for the overall experiment and specifically at the developmental timepoints 1 and 2 with the results function, using the Wald test at alpha = 0.05.

An annotation package was created for flowering dogwood using *AnnotationForge* 1.48 (Marc Carlson, 2025) and the EnTAP results. The proteome was annotated using the KEGG Automatic Annotation Server (KAAS) (Moriya *et al*., 2007). *ClusterProfiler* version 4.14.4 (Wu *et al*., 2021) was used for gene enrichment analysis using both annotation methods. To elucidate the identity of discovered MYB transcription factors (TFs), a phylogenetic tree was constructed with the neighbor-joining method using *clustalo* 1.2.4 (Sievers & Higgins, 2018), *trimAl* v1.4.rev15 (Capella-Gutiérrez *et al*., 2009), and *MEGA* version 11 (Tamura *et al*., 2021) (Table S3). One identified MYB was found to be truncated due to gene annotation error. To recover the full protein sequence, an in-house genome annotation pipeline was used, Ragnorak (https://github.com/ryandkuster/ragnarok).

### Contextualizing Results with Genomic Resources

The four chromosome-scale assemblies (‘Cherokee Brave’ Hap 1 and 2–pink-bracted and red-leafed, and ‘Appalachian Spring’ Hap 1 and 2–white-bracted and green-leafed) and their gene (from *BRAKER3*) and repeat (from *RepeatModeler*) annotations were visualized with *Persephone* (Persephone Software, LLC). Additionally, the BAM files of the HiFi reads and PAF files of the assemblies mapped back to the ‘Cherokee Brave’ Hap 2 genome were visualized within *Persephone*. The VCF file was visualized with *IGV* 2.13, and SNPs with diagnostic potential were noted. The regions flanking the group of diagnostic SNPs were inspected in *Persephone*, noting candidate genes relating to anthocyanin biosynthesis. For genes of interest, either from the list of differentially expressed genes, enrichment analysis, or visual inspection of the region and diagnostic SNPs, the normalized gene count throughout the experiment was calculated with *DESeq2*.

## Results

### Genome Assembly and Scaffolding

The haploid genome size of ‘Cherokee Brave’ was estimated by flow cytometry at 1.21 Gb. For both cultivars, PacBio HiFi and Hi-C data were generated. ‘Cherokee Brave’ was sequenced to 72× coverage with an average PacBio HiFi read length of 14,517 bp, whereas ‘Appalachian Spring’ was sequenced to 38× coverage with an average read length of 17,104 bp. Hi-C sequencing of ‘Appalachian Spring’ yielded 233 M read-pairs, of which 42.13% were same-strand high-quality and 35.64% were classified as informative read pairs. For ‘Cherokee Brave’, 343 M read-pairs were generated with 38.53% same-strand high-quality read pairs and 34.14% informative read pairs.

After assembly and scaffolding of the genome (Table S1), the final chromosome-level assembly for ‘Cherokee Brave’ had a genome size of 1,252.7 Mb with an N50 of 27.1 Mb for Haplotype 1 and a genome size of 1,253.0 Mb with an N50 of 47.4 Mb for Haplotype 2 (Table 1). The genome size of the final chromosome-level assembly for ‘Appalachian Spring’ was 1,266.7 Mb with an N50 of 17.2 Mb for Haplotype 1 and a genome size of 1,254.0 Mb and an N50 of 15.6 Mb for Haplotype 2 (Table 1 and Table S1). The total complete BUSCOs ranged from 99.1-99.3% across the four final assemblies (Table 1 and Table S1).

**Table 1.** Summary statistics of *Cornus florida ‘Cherokee Brave’* and ‘Appalachian Spring’ genome assembly and annotation annotations ranged between 98.0 and 98.2% for the four assemblies (Table S2). The percentage of genes that were functionally annotated by EnTAP ranged between 95.6 and 96.0% for the four assemblies (Table 1 and Table S2). The ratio of mono-exonic genes to multi-exonic genes ranged from 0.32 to 0.35 for the four assemblies (Table S2).

|  | Cherokee Brave' |  | Appalachian Spring' |  |
| --- | --- | --- | --- | --- |
|  | Hap 1 | Hap 2 | Hap 1 | Hap 2 |
| **Assembly Statistics** |  |  |  |  |
| Number of Scaffolds | 482 | 363 | 505 | 383 |
| Number of Contigs | 660 | 459 | 788 | 597 |
| Scaffold Total Length (Mb) | 1252.688 | 1253.041 | 1266.729 | 1254.111 |
| Contig Total Length (Mb) | 1252.599<br>0.007% gap | 1252.993<br>0.004% gap | 1266.588<br>0.011% gap | 1254.004<br>0.009% gap |
| Scaffold N/L50 (Mb) | 6/111.694 | 6/107.931 | 6/110.188 | 6/107.644 |
| Contig N/L50 (Mb) | 14/27.101 | 9/47.359 | 19/17.198 | 23/15.601 |
| Max scaffold length (Mb) | 140.531 | 140.401 | 140.746 | 140.819 |
| Max contig length (Mb) | 79.019 | 124.686 | 82.468 | 62.121 |
| Scaffolds > 50 KB (%) | 99.26 | 99.57 | 99.27 | 99.63 |
| Genome in chromosomes (%) | 96.8 | 97 | 96.3 | 96.4 |
| Complete BUSCOs (C) | 1602 (99.3%) | 1601 (99.2%) | 1603 (99.3%) | 1599 (99.1%) |
| Complete and single-copy BUSCOs (S) | 1528 (94.7%) | 1534 (95.0%) | 1523 (94.4%) | 1526 (94.6%) |
| **Annotation Statistics** |  |  |  |  |
| Number of genes | 28,695 | 28,558 | 28,768 | 28,615 |
| Number of mRNA | 32,300 | 32,145 | 32,341 | 32,105 |
| EnTAP annotation rate (%) | 95.81 | 95.84 | 95.99 | 96 |

We visualized the genome assemblies using multiple methods throughout the assembly process to qualitatively assess the genomes. The combination of high intensity interaction along the diagonal and small amounts of debris (i.e., unplaced scaffolds and contigs) within the Hi-C contact map indicates high-quality genomes (Figure S2). Between 99.3% and 99.6% of loci within the P25×P28 linkage map from Moreau et al. (2022) were anchored into the equivalent chromosome of our assemblies (Figure S3). SyRI and plotsr were used as the final visualization to qualitatively assess genome quality (Figure S4).

### Genome Annotation

To prepare for genome annotation, the repeats within the scaffolded assemblies were masked, resulting in 71.8% of both the diploid ‘Cherokee Brave’ and ‘Appalachian Spring’ assemblies being soft-masked (breakdown of repeat classes in Figure S5; Table S2). For ‘Cherokee Brave’, 28,695 genes were identified for Hap 1 and 28,558 for Hap 2 (Table 1 and Table S2). For ‘Appalachian Spring’, 28,768 genes were annotated for Hap 1 and 28,615 genes for Hap 2 (Table 1 and Table S2). BUSCO scores for genome

### Population Genotyping

GBS was used to genotype the “pseudo-F2” population of 125 individuals. The number of sequences per sample ranged from 5.6 to 16 M before filtering and from 5.2 to 16 M after filtering with *fastp*. One sample failed sequencing and had ∼20k reads (F2-06-65). After combining the three sequencing runs for the parents (97-6 and 97-7), they had a total of 21.0 M and 18.2 M reads, respectively. After filtering, they yielded 20.0 M and 17.8 M sequences, respectively. Due to ‘Cherokee Brave’ Hap 2 having the highest contig N50 and lowest number of contigs, it was selected as the reference genome for all analyses. The average mapping rate was 98.0% across all samples, and the resulting BAM files were used in the GATK pipeline to call SNPs. Directly out of GATK, there were 1,315,679 SNPs, and after filtering according to GATK’s best practices guidelines, there were 1,172,327 SNPs. A parameter sweep of additional filtering was performed for maximum missing genotypes (5-20%), minimum read depth (10), and minor allele frequency (0.02-0.05), resulting in the number of SNPs ranging from 7,566 to 12,646. To ensure there was no underlying structure in the SNPs in general or at the varying missing data levels, the output from the varying filtering options was visualized via PCA (Figure S6a). No structure was observed in any iteration. A kinship matrix was calculated and visualized as a heatmap to identify any unrelated individuals in the “pseudo-F2” population. Eight individuals had an average negative kinship value (i.e., brown in the heatmap) and were removed from the dataset (Figure S6b). The VCF filtered for 20% maximum missing and 0.05 minor allele frequency was used for downstream applications.

### Linkage Map Construction

The 116 individuals and 12,646 SNPs left after filtering were used for linkage map construction using both the recombination and genomic position information. A total of 2,747 SNPs were removed due to missing data in the parents, and 5,440 uninformative SNPs were removed because the parents were homozygous at these loci. An additional 2,307 SNPs were removed because missing data was greater than 10%. From here, the SNPs were binned to remove redundant markers, with an average of 1.76 markers/bin, resulting in 2,587 non-redundant markers (removed 1,962 SNPs). Markers were dropped if they had significant segregation distortion or if they had an unexpected pattern in the recombination matrix. The final linkage map contained 1,148 markers, with 792 segregating 1:2:1 (ab × ab), 210 segregating 1:1 (ab × aa), and 145 segregating 1:1 (aa × ab). The average distance between markers is 0.26 cM, and the total map spans 2,449 cM (Figure 2 and Table 2).

**Figure 2.**
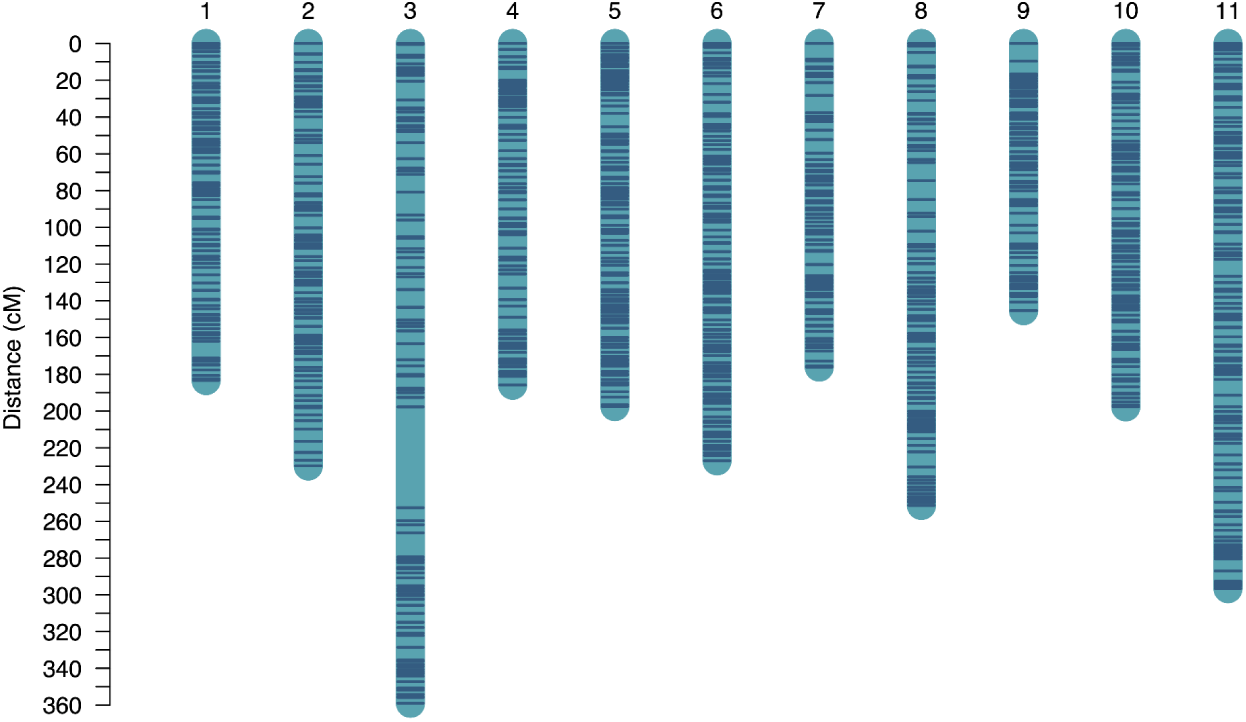
Linkage map constructed from *Cornus florida* ‘Appalachian Spring’ × ‘Cherokee Brave’ “pseudo-F2” population. Includes 1148 markers with an average distance of 0.26 cM between markers.

**Table 2.** Summary statistics for linkage map constructed from “pseudo-F2” population of *Cornus florida*.

| LG | Number of Markers | Total Distance (cM) | Average Distance (cM) | Maximum Distance (cM) |
| --- | --- | --- | --- | --- |
| 1 | 100 | 183.3 | 1.9 | 9.2 |
| 2 | 102 | 229.8 | 2.3 | 7.3 |
| 3 | 97 | 359 | 3.7 | 54.8 |
| 4 | 83 | 185.8 | 2.3 | 7.7 |
| 5 | 118 | 197.5 | 1.7 | 7.3 |
| 6 | 123 | 227 | 1.9 | 6.3 |
| 7 | 86 | 176 | 2.1 | 9.5 |
| 8 | 110 | 251.3 | 2.3 | 10.3 |
| 9 | 84 | 145.3 | 1.8 | 9.6 |
| 10 | 103 | 197.6 | 1.9 | 5.9 |
| 11 | 142 | 296.6 | 2.1 | 9.2 |
| Total | 1148 | 2449.2 | 2.2 | 54.8 |

### Population Phenotyping

To find the most effective method to identify QTL associated with bract color, multiple traits were measured for the bracts and compared to categorical methods. Categorical data with five classes was collected for both the leaves and bracts: 1 = white bract/green leaf, 2 = less than half pink bract/red leaf, 3 = half pink bract/red leaf, 4 = more than half pink bract/red leaf, 5 = almost completely pink bract/red leaf (see Figure 1b for visual representation). To better capture the variation of bract color in the population, we also took quantitative measurements, including L*a*b* values from a colorimeter and L*a*b* and RGB values from images. For the categorical data, all individuals classified with bract color (category ≥ 2) were also classified as having leaf color (white bracted/green leafed = 84, pink bracted/red leafed = 24). Out of the eight trees that did not flower during the data collection year, six were red-leaved. For the five-class system, trees were not consistently scored as the same color intensity for leaves and bracts (96/108 match; for distributions, see Figure 3a). As 50 of the original 175 trees have died, segregation ratios calculated based on these phenotype frequencies are not expected to be accurate.

**Figure 3.**
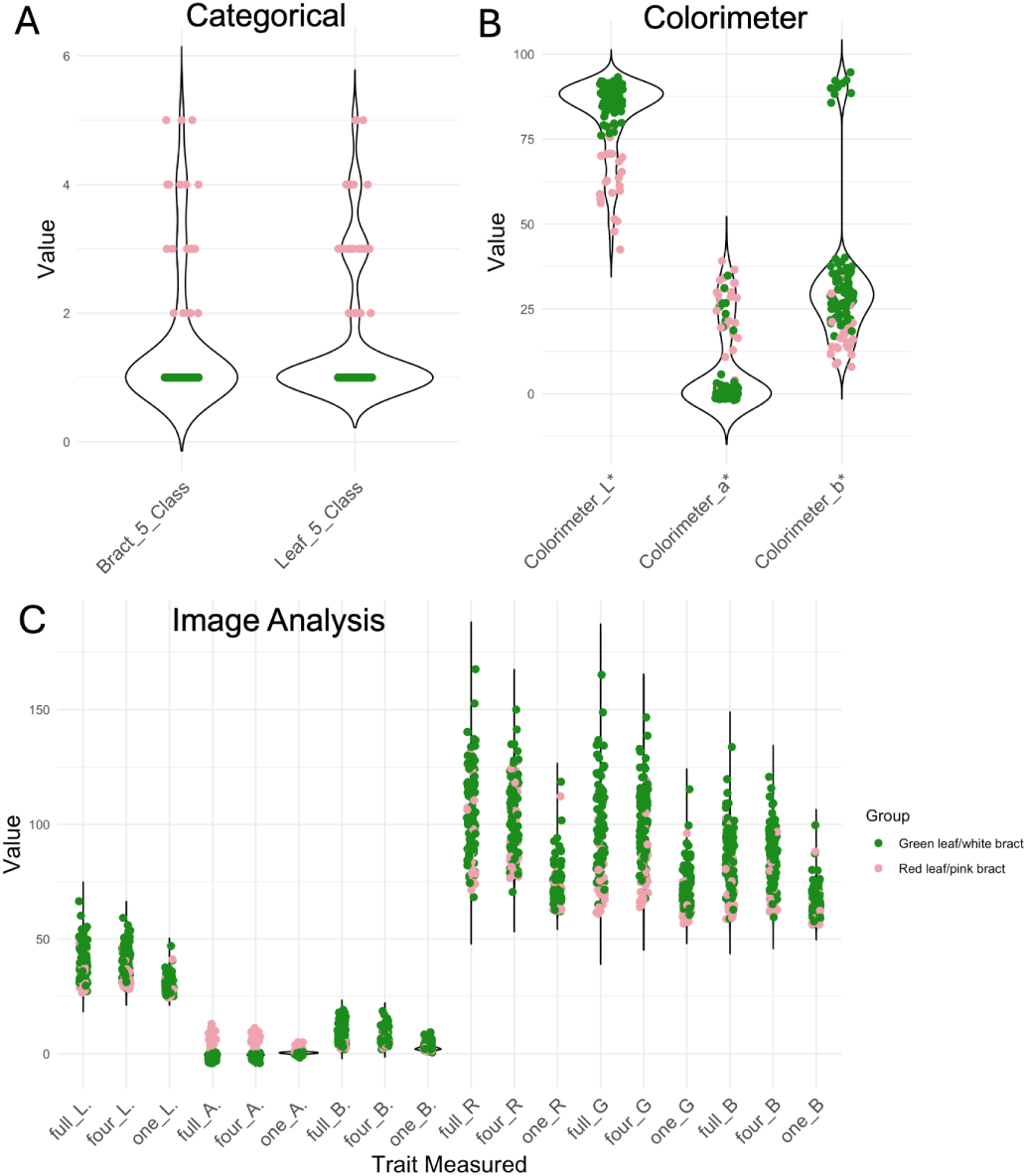
Distributions of phenotypes measured of *Cornus florida* ‘Cherokee Brave’ × ‘Appalachian Spring’. A) categorical data B) average colorimeter L*a*b* readings for each individual C) average L*a*b* and RGB values from segmented images of the full inflorescence, four bracts pulled off the main inflorescence, and one bract. Green dots are individuals classified as white bracted/green leafed based on categorical data (n=84) and pink dots are individuals classified as pink bracted/red leafed (n=24).

A total of 1,044 pictures and 349 colorimeter readings were taken of inflorescences of the 108 individuals. Three individuals were missing a full inflorescence picture, and one individual only had one inflorescence on the tree, and therefore only had one colorimeter reading and one image for each picture type. All other individuals had three data points per metric. The distributions of the average L*a*b* color values for an individual from the colorimeter are available in Figure 3b. The distributions of the average L*a*b* and RGB values for an individual from the three image types are available in Figure 3c. The only trait measurements that clearly separated the pigmented trees from the non-pigmented trees in this figure were the a* value from image analysis and the L* value from the colorimeter.

### Anthocyanin Profiling

Anthocyanin profiling was conducted on ‘Cherokee Brave’, the pink-bracted, red-leafed parent of the “pseudo-F2” population. The primary anthocyanins identified in the leaves and bracts during profiling were Delphinidin 3-O-galactoside and Delphinidin 3-O-sambubioside (Figure S7). The following were also detected in the leaves and bracts: Cyanidin, Cyanidin 3-O-2G-glucosylrutinoside, Cyanidin 3-O-galactoside, Cyanidin 3-O-lathyroside, Cyanidin 3,5-O-diglucoside, Delphinidin, Delphinidin 3-O-arabinoside, Delphinidin 3,5-O-diglucoside, Malvidin 3-O-(6’’-malonylglucoside), Malvidin 3-O-arabinoside, Pelargonidin, Pelargonidin 3-O-galactoside, Peonidin 3,5-O-diglucoside, Petunidin, Petunidin 3-O-galactoside, Petunidin 3,5-O-diglucoside, and Quercetin 3-rutinoside(Rutin). The following were only detected in the leaves: Cyanidin 3-O-rutinoside, Cyanidin 3-O-sophoroside 5-O-glucoside, and Malvidin. The following were tested for but not detected in either of our samples: Malvidin 3-O-galactoside, Peonidin.

### QTL Mapping

CIM was used for all trait measurements to test phenotyping methods for future bract quantitative studies. A single high-effect QTL was consistently detected on LG 9 for 12/23 traits, primarily the L* (lightness), a* (red-green color axis) values, and categorical data (Figure S8a). The bract locus with the highest LOD score (Figure 4) was from the four-bract image a* (i.e., the image with the bracts pulled off the main inflorescence). Permutation thresholds were calculated for the four-bract image, colorimeter a*, bract categorical, and leaf categorical (Figure S8c), and only the LOD peak for colorimeter a* is below the threshold. The leaf locus had the highest LOD peak of any of the methods and had a smaller locus than the other methods. The four-bract a* locus is a 14Mb locus flanked by markers Chr09_38583748 and Chr09_52556621. The leaf locus is smaller, flanked by markers Chr09_38583748 and Chr09_46098776 (i.e., 7.5Mb locus). The marker coverage in this area is sparse, therefore, the four-bract a* locus was used as the locus boundaries in downstream analyses. There are 209 genes in this region based on the ‘Cherokee Brave’ Hap 2 genome annotation.

**Figure 4.**
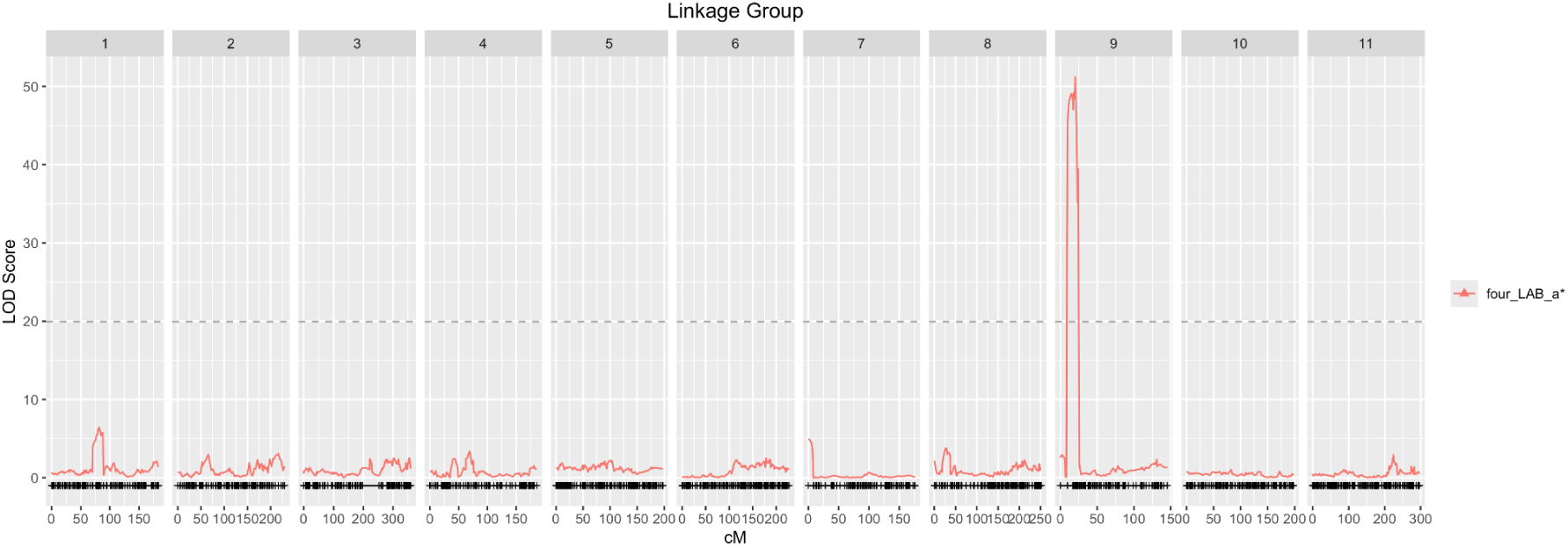
*Cornus florida* ‘Appalachian Spring’ × ‘Cherokee Brave’ “pseudo-F2” population QTL results for bract color. Bract color was measured from an image of the four bracts pulled off the main inflorescence. The image was segmented to just the bracts and the segmented pixels were used to calculate the average a* (red-green color axis in L*a*b* colorscale) across the segmented pixels. The threshold was calculated using 1000 permutations with alpha = 0.05.

### Differential Gene Expression

The differential expression analysis was conducted using bracts and leaves on a developmental time course of ‘Appalachian Spring’ and ‘Cherokee Brave’. Throughout the three developmental timepoints, a total of 6,021 and 11,645 genes were differentially expressed in the bracts and leaves, respectively. Wald tests were conducted for Timepoint 1 (T1) and Timepoint 2 (T2) because color appears early on in ‘Cherokee Brave’ as it is developing. For T1, a total of 932 and 3863 genes were upregulated and 1,376 and 3,667 genes were downregulated in ‘Cherokee Brave’ as compared to ‘Appalachian Spring’ in the leaves and bracts, respectively. For T2, a total of 3,091 and 1,899 genes were upregulated and 3,351 and 1,917 genes were downregulated in ‘Cherokee Brave’ as compared to ‘Appalachian Spring’ in the leaves and bracts, respectively.

Out of the 209 genes within the 14Mb locus, there were 59 differentially expressed genes (DEGs) in T1 for bracts, 20 in T1 for leaves, 42 in T2 for bracts, and 28 in T2 for leaves (Wald Test at alpha = 0.05). Out of these DEGs, there are 10 common DEGs at T1 for the leaves and bracts, 19 for T2, and 15 for the whole experiment (Table S3-S5). The functions of these genes were investigated using EnTAP functional assignments (Table S3-S5). One of the DEGs in T1 and T2 has a MYB DNA-binding domain (*g19533*). Expression levels of *g19533* are consistently higher in ‘Cherokee Brave’ than ‘Appalachian Spring’ throughout the experiment (Figure 5). The protein sequence of g19533 was compared to other plant MYB TFs and found to be truncated in length due to a gene annotation error. Using an in-house annotation pipeline, Ragnorak (https://github.com/ryandkuster/ragnarok), Chr09 was annotated, and the full protein sequence was recovered. In the phylogenetic tree for the gene family, the full protein coding sequence is grouped with a maize MYB-related protein (P27898.1) with a bootstrap support of 60. No functions have been assigned to this MYB in maize. Other notable DEGs in the QTL region include three with RING finger binding domains (*g19497*, *g19556,* and *g19590*). Of these, *g19497* (a RING zinc finger-C3HC4) and *g19590* have consistent patterns of expression between the leaves and bracts for the entire experiment, whereas *g19556* has the same pattern for T1 and T2, but not T3 (Figure 5).

**Figure 5.**
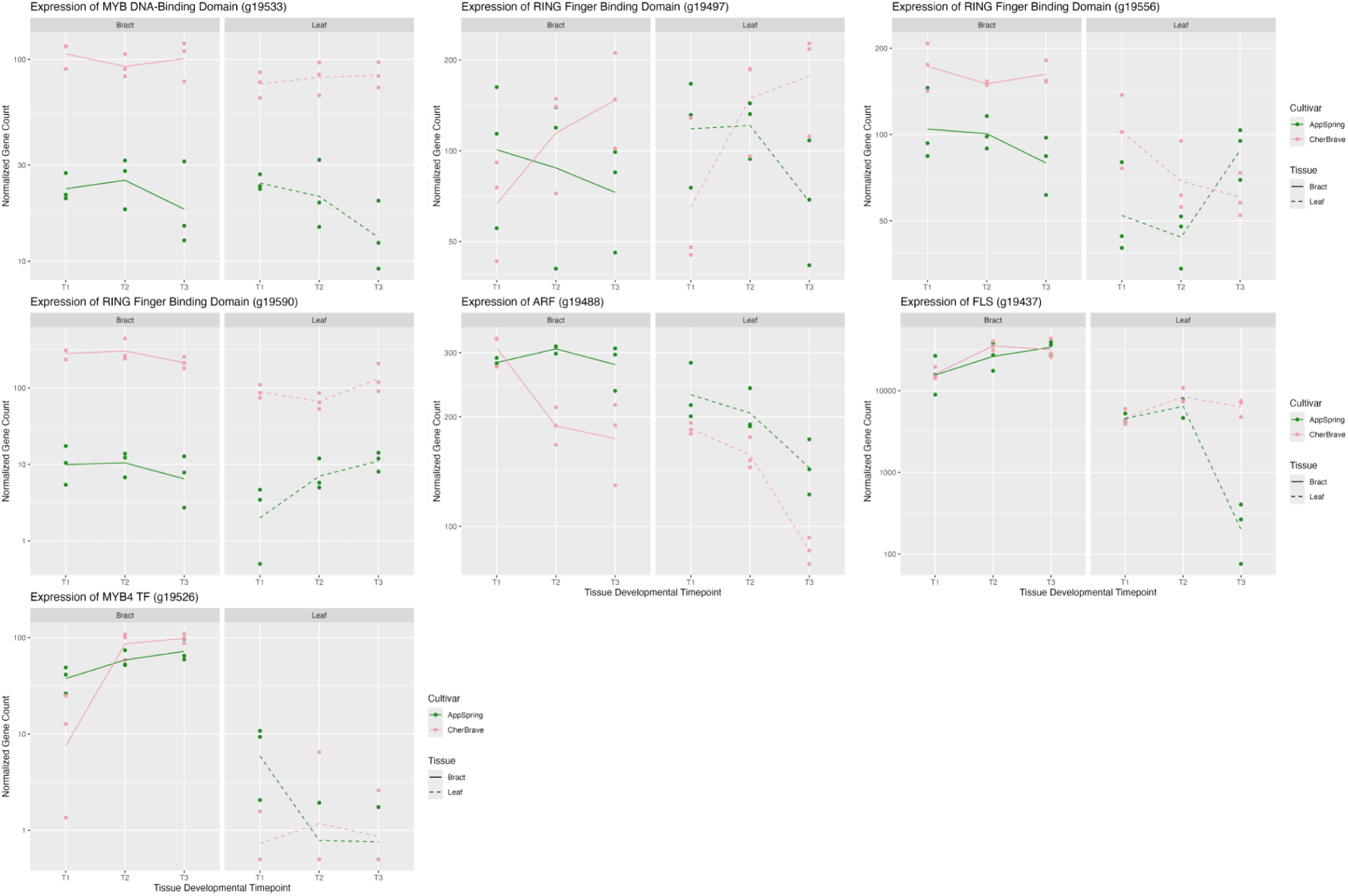
Expression levels of seven genes of interest in *Cornus florida* ‘Cherokee Brave’ and ‘Appalachian Spring’ leaves and bracts across three developmental timepoints. Timepoints abbreviated as: T1 = breaking bud, T2 = expanding tissue, T3 = fully expanded tissue. ‘Appalachian Spring’ has green leaves, white bracts, and gene counts are in green. ‘Cherokee Brave’ has red leaves, pink bracts, and gene counts are in pink. Gene counts in the bracts have a solid line and gene counts in the leaves have a dashed line.

Another independent functional assignment tool, KEGG Automatic Annotation Server (KAAS)was used to assign KEGG Orthology (KO) IDs to all ‘Cherokee Brave’ Hap2 genes. Only 35.6% of all genes were assigned a KO. Within the bract locus, only 27.8% of the genes were assigned a KO ID. KO assignments were inspected for phenylpropanoid biosynthesis, flavonoid biosynthesis, anthocyanin biosynthesis, or plant hormone signal transduction pathway involvement. Through this method, an *Auxin Response Factor* (*ARF*; *g19488*) and an array of *Flavonol Synthase* (*FLS*; *g19433*, *g19434, g19436, g19437*) genes were identified. These genes are not differentially expressed at T1 and do not have consistent patterns of expression in the leaves and bracts (Figure 5 and Figure S9). Fewer than 100 transcripts/rep were counted for *g19433*, *g19434, g19436* (Figure S9), and more than 1000 transcripts/rep were counted for *g19437* (Figure 5).

Additionally, the maximum likelihood estimates of the log2fold expression changes of the genes were used to identify genes of interest. By screening log2fold changes < −1.5 in T1 in the leaves and bracts, a MYB transcription factor was identified (*g19526*). The expression was higher in ‘Appalachian Spring’ than ‘Cherokee Brave’ at T1 (Figure 5). This gene was identified as *MYB14* by EnTAP, however, when blasted within the UniProt database, it was most similar to *MYB4*. In the neighbor-joining phylogenetic tree, *g19526* grouped with *Arabidopsis thaliana AT4G38620 MYB4* (Figure S10; Table S6).

GO and KEGG enrichment analysis was performed for the overall time course experiment. In both the leaves and bracts in T1, there were several GO terms relating to hormone detection and response (Figure S11). In the leaves, detection of hormone stimuli was activated at T1 in ‘Cherokee Brave’ and not picked up in the overall experiment. In the bracts, responses to hormone stimuli were suppressed at T1 in ‘Cherokee Brave’ but activated overall throughout the experiment. Cultivar-specific differences in expression of anthocyanin-related genes were visualized within the KEGG pathways for phenylpropanoid biosynthesis (ko00940), flavonoid biosynthesis (ko00941), and anthocyanin biosynthesis (ko00942). Results for bracts and leaves are located on the same plot (Figures S12-S17).

### Contextualizing Results with Genomic Resources

Upon inspection of the “pseudo-F2” VCF file, within the bract locus, there is a 274,961 basepair window (40907723-41182684) where the SNPs predict the presence/absence of bract and leaf color (0/0 = pink bracts, 0/1 = white or slightly blushed bracts, and 1/1 = white bracts). The only exceptions are due to genotypic missing data. In the HiFi reads mapped back to the ‘Cherokee Brave’ genome, these SNPs in ‘Cherokee Brave’ (pink-bracted) are 0/0, and in ‘Appalachian Spring’ (white-bracted) are 1/1. Using this information, combined with the biological knowledge of needing two generations to recover reliable pink bracts, we searched for variations within and around our genes of interest with the homozygous reference in ‘Cherokee Brave’ and the homozygous variant in ‘Appalachian Spring’.

For the DEGs, three of them have variants with the expected pattern. The *g19533* gene (contains MYB DNA-binding domain) has two copies in ‘Cherokee Brave’ and zero in ‘Appalachian Spring’, and within ‘Cherokee Brave’, there are two isoforms in Chromosome 9; where 55 bp upstream of *g19497* (contains RING Zinc Finger C3HC4-type binding domain) is a 7 bp indel within a microsatellite region with the expected pattern. The *g19556* gene (RING finger binding domain) has three nonsynonymous SNPs and one synonymous SNP in the expected genotypic pattern, among others in the introns. However all three of these genes are outside of the diagnostic SNP window.

In the diagnostic SNP locus, there are some genes of interest with the same pattern of SNPs. Other genes of interest include an array of *Walls are Thin 1* (*WAT1*; *g19420*-*g19422*) and a gene with a RING Zinc Finger C3HC4-type binding domain (*g19416*). All genes have homozygous reference SNPs in ‘Cherokee Brave’ and homozygous alternate SNPs in ‘Appalachian Spring’. These genes are not differentially expressed within our time course experiment (Figure S10). A summary of genes of interest and corresponding evidence can be found in Figure 6.

**Figure 6.**
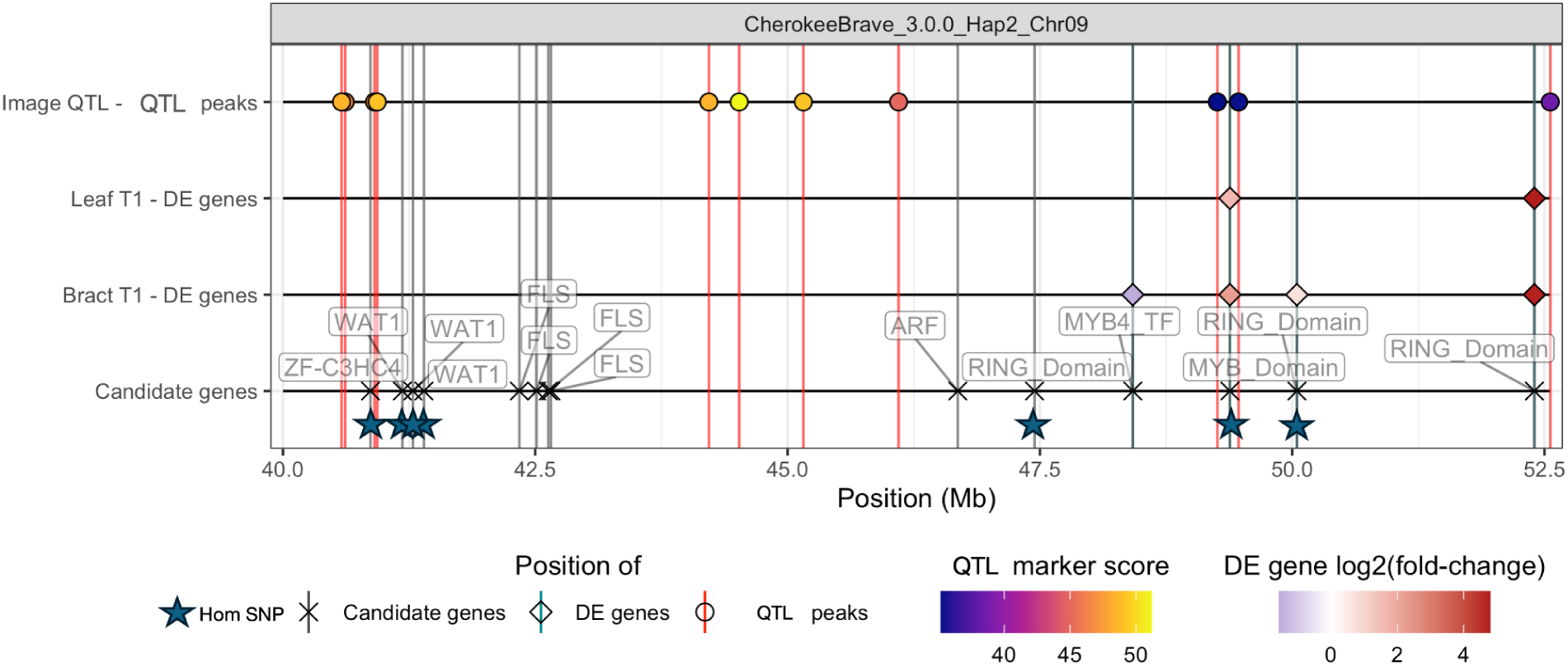
Hidecan plot of genes of interest within QTL. The first line contains QTL LOD scores for the four-bract a* value from image analysis. The second line contains significant (alpha = 0.05) differentially expressed genes at timepoint 1 in the leaves. The third line contains significant (alpha = 0.05) differentially expressed genes at timepoint 1 in the bracts. Identified candidate genes are located on the fourth line. Blue stars indicate genes with homozygous reference variants in *Cornus florida* ‘Cherokee Brave’ (pink-bracted) and homozygous alternate variants in ‘Appalachian Spring’.

## Discussion

As an ornamental tree with limited genomic resources, breeding for flowering dogwood has posed several challenges. The long 3-to-5 year generation time combined with the space requirements of growing large progeny of trees have hindered progress at the only public breeding program for flowering dogwoods in the U.S., the Rutgers University Woody Ornamentals Breeding Program (Molnar, 2022). The space, resource requirements, and time burden associated with phenotyping and caring for trees without desired phenotypes would be decreased with a marker-assisted-selection (MAS) based breeding program where individuals could be screened in the greenhouse before field-planting. Additionally, if the genetics of traits of interest were better understood, the traits could be combined in cultivars and released to consumers faster. Within the consumer market, dogwoods annually generate more than $31 million (USD) in wholesale and retail sales from more than 1 million trees in the U.S. (USDA-NASS, 2020). This market share could grow if novel trait combinations desired by consumers were more swiftly released.

Developing high-quality genomic resources serves as the backbone for incorporating MAS into a historically observational-based breeding strategy. These resources have greatly impacted breeding in other non-model plants, like strawberry (Kim *et al*., 2025), flowering cherry (Nie *et al*., 2023), and rose (Zhang *et al*., 2024). In this study, we present the first high-quality, chromosome-scale, diploid, annotated assemblies for flowering dogwood. We assembled and annotated assemblies for *C. florida* ‘Cherokee Brave’, a pink-bracted and red-leafed cultivar, and ‘Appalachian Spring’, a white-bracted and green-leafed cultivar. Our four haplotype assemblies ranged in size from 1,252.688 Mb to 1,266.729 Mb. This is close to our ‘Cherokee Brave’ flow cytometry estimate of 1,210 Mb, but smaller than the previously reported genome size of 1,500 Mb for ‘Cherokee Brave’ and 1,560 Mb for ‘Appalachian Spring’ (Shearer & Ranney, 2013).

This study focused on using the developed genomic resources to gain insight into the mechanisms underlying bract and leaf color in flowering dogwood. To accomplish this, we leveraged a “pseudo-F2” population generated from ‘Cherokee Brave’ and ‘Appalachian Spring’. ‘Cherokee Brave’ and ‘Appalachian Spring’ were crossed to obtain an F1 generation, and then two F1s (97-7 and 97-6) were crossed to produce the “pseudo-F2” population. The “pseudo-F2” population was used for linkage mapping, and the P1 generation (‘Cherokee Brave’ and ‘Appalachian Spring’) was used for identification of differentially expressed genes within the locus.

To date, the only QTL mapping study involving flowering dogwood color used an SSR-based linkage map (Wang *et al*., 2009) and investigated only the leaf color of seedlings (Wadl *et al*., 2011). Four putative single-marker QTL were identified by Wadl et al. (2011) on LG3, 6, and 8 (LG10, 6, 5, respectively in our map and genome). However, the QTL explained a low percentage of phenotypic variance, and QTL instability was detected. High-density linkage mapping has been instrumental in identifying candidate genes in chrysanthemum flower color (Song *et al*., 2023) and grape berry color (Sun *et al*., 2020). Therefore, a new, high-density linkage map and additional phenotyping methods were applied to this study.

A total of 12/23 traits, primarily the L* (lightness), a* (red-green color axis) values, and categorical data, detected a single high-effect locus on LG9 associated with bract and leaf color. SNPs within this locus can predict the presence or absence of bract and leaf coloration with perfect accuracy within our population. Homozygous reference (0/0) reliably indicates pink bracts and red leaves, whereas homozygous alternate (1/1) reliably indicates white bracts and green leaves. In contrast, the heterozygotes (0/1) usually indicate white bracts and green leaves but can display some blush pigmentation if there is cold damage. These conditions are consistent with prior knowledge of F1 crosses between white-bracted individuals and pink-bracted individuals. Considering the homozygous reference SNPs within LG9 do not differentiate the category 2-to-5 colorations in the segregating population, additional work is required to pinpoint modifier loci that regulate the intensity of the pigmentation.

This study also addressed best practices for phenotyping bract color for future research. The five-class leaf categorical data had the highest LOD score, followed by the a* value from the four-bract image analysis. Both peaks were above the permutation threshold for these traits. Eight individuals were not flowering the year the population was phenotyped, therefore, the leaf five-class categorical data likely had the highest LOD score because there were the most individuals included for that trait (n=116 vs. n=108). From an implementation standpoint, the easiest and most straightforward phenotyping method is classifying individuals into 5 categories. However, it should be noted that if the five-class leaf locus would have been used, we would have missed the identification of all of our differentially expressed genes with the expected variant patterns due to the leaf locus being smaller. This could be a reflection of our sparse linkage map in this area, the imprecision of the categorical data, or separate but close loci for the leaves and bracts.

The next easiest method to implement was the colorimeter, but the LOD peak was below the permutation threshold. There were nine white-bracted individuals with a* values consistent with category 3-to-5 pink bracts, which likely impacted the permutation threshold. This inconsistency was likely due to human error with either the recording of values or the usage of the colorimeter, as the a* values from image analysis were close to 0, as expected. Studies in groundcover rose (*Rosa* × *hybrida*) (Schmitzer *et al*., 2010) and althea (*Hibiscus syriacus*) (Lattier & Contreras, 2020) have had success using colorimeters in breeding programs. From the practical standpoint of managing a breeding program, we recommend supplementing categorical data with colorimeter data and placing checks through the data recording process. Other association mapping studies have also benefited from quantifying a single trait with multiple metrics (Shu *et al*., 2024). Therefore, recording both colorimeter and categorical data could help identify new loci. The process of capturing images and image processing is time-consuming, but could be worthwhile to pursue with small populations to increase detection power.

As expected with a small “pseudo-F2” population, the identified locus was large at 14Mb with 209 genes. With the complicated regulatory network of anthocyanin biosynthesis, there is vast diversity in methods of activation and repression of the pathway (for a review of known regulatory elements within a non-model species, see: Chen *et al*., 2021). While activation of anthocyanin production is highly conserved with the WDR complex, there is great diversity in the methods of repression, including direct repression, protein-protein interaction, and post-transcriptional gene silencing (LaFountain & Yuan, 2021). Within our locus, there are several candidate genes related to the anthocyanin biosynthetic pathway. Broadly, the candidate genes within the locus encode structural anthocyanin enzymes, MYB binding domains, RING-binding domains, and auxin-response proteins.

To narrow down candidate genes, a time-course RNAseq experiment using the P1 generation (‘Cherokee Brave’ and ‘Appalachian Spring’) was conducted. The most promising candidate genes are *g19533*, with a MYB-DNA Binding domain, and *g19556*, with a RING finger binding domain. MYB TFs are some of the most well-known variables for regulating anthocyanin pigmentation, and commonly increase anthocyanin accumulation (Li *et al*., 2016; Zhang *et al*., 2024). Many regulatory elements can impact the transcription of a gene or the function of a final protein. RING finger binding domains are commonly indicative of E3 ligases, which have been demonstrated to mark MYB TF for degradation (An *et al*., 2017, 2020). Proteins with RING finger binding domains have also been demonstrated to stabilize MYB TF (Yang *et al*., 2024). In this experiment, expression levels of *g19533* were consistently higher in ‘Cherokee Brave’ than ‘Appalachian Spring’ throughout all timepoints, suggesting it could be an activating TF. Within Chromosome 9 of the genome assemblies, *g19533* is present in both ‘Cherokee Brave’ Hap 1 and 2, however, no blast hits are present within either ‘Appalachian Spring’ Chromosome 9 haplotype assemblies. There is a 5,241 bp homozygous reference insertion in ‘Cherokee Brave’ and a homozygous alternate deletion in ‘Appalachian Spring’ that contains this gene. There is spurious RNAseq read mapping occurring from ‘Appalachian Spring’ to this gene, despite the clear evidence of a deletion from the assemblies and HiFi data. These spurious reads mapping to *g19533*, map exactly to a homolog on ‘Appalachian Spring’ Chr08 Hap 2. Based on the neighbor-joining tree, this MYB TF grouped with a maize MYB TF of unknown function. However, the bootstrap support was low at 60, therefore, additional research is needed to identify this MYB TF and its exact function. For *g19556,* significant expression differences between ‘Cherokee Brave’ and ‘Appalachian Spring’ at T1 were only present in the bracts, however, the trend of higher expression in ‘Cherokee Brave’ than ‘Appalachian Spring’ was present in the leaves as well. Genomic variation within this gene also matches our expected pattern, with SNPs following the expected segregation pattern in the first two exons of the gene, among others in the introns.

In this study, we leveraged our genomic resources to identify many differentially expressed and anthocyanin-related genes in the locus of interest. However, there were only two with both differential expression evidence and variants with the expected homozygous reference in ‘Cherokee Brave’ and homozygous alternate in ‘Appalachian Spring’. Both genes, *g19533*, with a MYB-DNA Binding domain, and *g19556*, with a RING finger binding domain, could play a role in regulating the phenotypic differences observed within the bracts and leaves of ‘Cherokee Brave’ and ‘Appalachian Spring’. Linkage mapping studies in other genetic backgrounds could be used to confirm and build upon the results presented here.

## Conclusions

We established genomic resources for flowering dogwood and implemented them in identifying genes of interest involved in the anthocyanin pigmentation accumulation in ‘Cherokee Brave’. The identified genes with a MYB binding domain and RING finger binding domain are promising candidates for the binary trait of whether or not anthocyanin pigmentation accumulation occurs. Additional research is necessary to pinpoint modifier loci that contribute to the gradients of bract and leaf color variation observed within the segregating population. These genomic resources, candidate genes, and associated SNPs will be used in dogwood breeding efforts to speed up combining traits of interest for consumers.

## Acknowledgements

The authors would like to thank Holly Brabazon, Allyson Dekovich, Beant Kapoor, Lav Yadav, and Matthew Huff for assisting with phenotyping the “pseudo-F2” population. The authors would also like to thank Jon Beever and the UTIA Genomics Center for library prep and sequencing of part of the timecourse RNAseq experiment.

## Author Contributions

Design of the research: M.E.S., R.N.T., T.P.H., D.H., D.S., W.E.K, M.N., T.J.M; performance of the research: T.P.H., S.L.B., J.L-M, A.H., S.A.S., R.C.Y., A.H-K., B.E.S.; data analysis, collection, or interpretation: T.P.H. and writing the manuscript: T.P.H. All authors critically reviewed and approved the manuscript.

## Data Availability

All ‘Cherokee Brave’ data is available under the NCBI Umbrella Project PRJNA989662, which include ‘Cherokee Brave’ Hap 1 (PRJNA988530) and Hap 2 (PRJNA988529) annotated genome assemblies. Data was uploaded to Hap 2 (BioProject PRJNA988529), and includes the raw PacBio HiFi, Hi-C, timecourse RNAseq, and demultiplexed GBS reads. All ‘Appalachian Spring’ data is also under the ‘Cherokee Brave’ Hap 2 BioProject (PRJNA988529). Custom scripts used in this study are available https://github.com/trinityhamm/Flowering-Dogwood-Color. Additional datasets and output are available from Ag Data Commons doi: (waiting on DOI).

## Supporting Information

Figure S1. Image examples for image-based phenotyping

Figure S2. Hi-C Contact maps for the four assemblies

Figure S3. Linkage Map vs. Genome assemblies for the four assemblies

Figure S4. SyRI plots of Hap 1 vs. Hap 2

Figure S5. Repeat classes masked in assemblies

Figure S6. PCA of filtered SNPs and kinship matrix of “pseudo-F2”

Figure S7. Anthocyanin profiling results of ‘Cherokee Brave’

Figure S8. QTL results

Figure S9. Expression of genes of interest

Figure S10. Neighbor-joining phylogenetic tree of MYB transcription factors

Figure S11. Activated and suppressed enriched GO terms

Figure S12. Log2fold change of KEGG phenylpropanoid biosynthetic pathway T1

Figure S13. Log2fold change of KEGG phenylpropanoid biosynthetic pathway T2

Figure S14. Log2fold change of KEGG flavonoid biosynthetic pathway T1

Figure S15. Log2fold change of KEGG flavonoid biosynthetic pathway T2

Figure S16. Log2fold change of KEGG anthocyanin biosynthetic pathway T1

Figure S17. Log2fold change of KEGG anthocyanin biosynthetic pathway T2

Methods S1. Supplementary methods

Table S1. Genome assembly statistics

Table S2. Genome annotation statistics

Table S3. EnTAP results for T1 differentially expressed genes

Table S4. EnTAP results for T2 differentially expressed genes

Table S5. EnTAP results for differentially expressed genes across the entire experiment

Table S6. MYB accession numbers used for neighbor-joining tree

