## Supplementary Figures and Methods for "Chromosome-Scale Assemblies of Flowering Dogwood Cultivars Enable Identification of Candidate Genes Regulating Anthocyanin Biosynthesis in Leaves and Bracts"

The following Supporting Information is available for this article:

Figure S1. Image examples for image-based phenotyping
Figure S2. Hi-C Contact maps for the four assemblies
Figure S3. Linkage Map vs. Genome assemblies for the four assemblies
Figure S4. SyRI plots of Hap 1 vs. Hap 2
Figure S5. Repeat classes masked in assemblies
Figure S6. PCA of filtered SNPs and kinship matrix of "pseudo-F2"
Figure S7. Anthocyanin profiling results of 'Cherokee Brave'
Figure S8. QTL results
Figure S9. Expression of genes of interest
Figure S10. Neighbor-joining phylogenetic tree of MYB transcription factors
Figure S11. Activated and suppressed enriched GO terms
Figure S12. Log2fold change of KEGG phenylpropanoid biosynthetic pathway T1
Figure S13. Log2fold change of KEGG phenylpropanoid biosynthetic pathway T2
Figure S14. Log2fold change of KEGG flavonoid biosynthetic pathway T1
Figure S15. Log2fold change of KEGG flavonoid biosynthetic pathway T2
Figure S16. Log2fold change of KEGG anthocyanin biosynthetic pathway T1
Figure S17. Log2fold change of KEGG anthocyanin biosynthetic pathway T2
Methods S1. Supplementary methods
Table S1. Genome assembly statistics
Table S2. Genome annotation statistics
Table S3. EnTAP results for T1 differentially expressed genes
Table S4. EnTAP results for T2 differentially expressed genes
Table S5. EnTAP results for differentially expressed genes across the entire experiment
Table S6. MYB accession numbers used for neighbor-joining tree

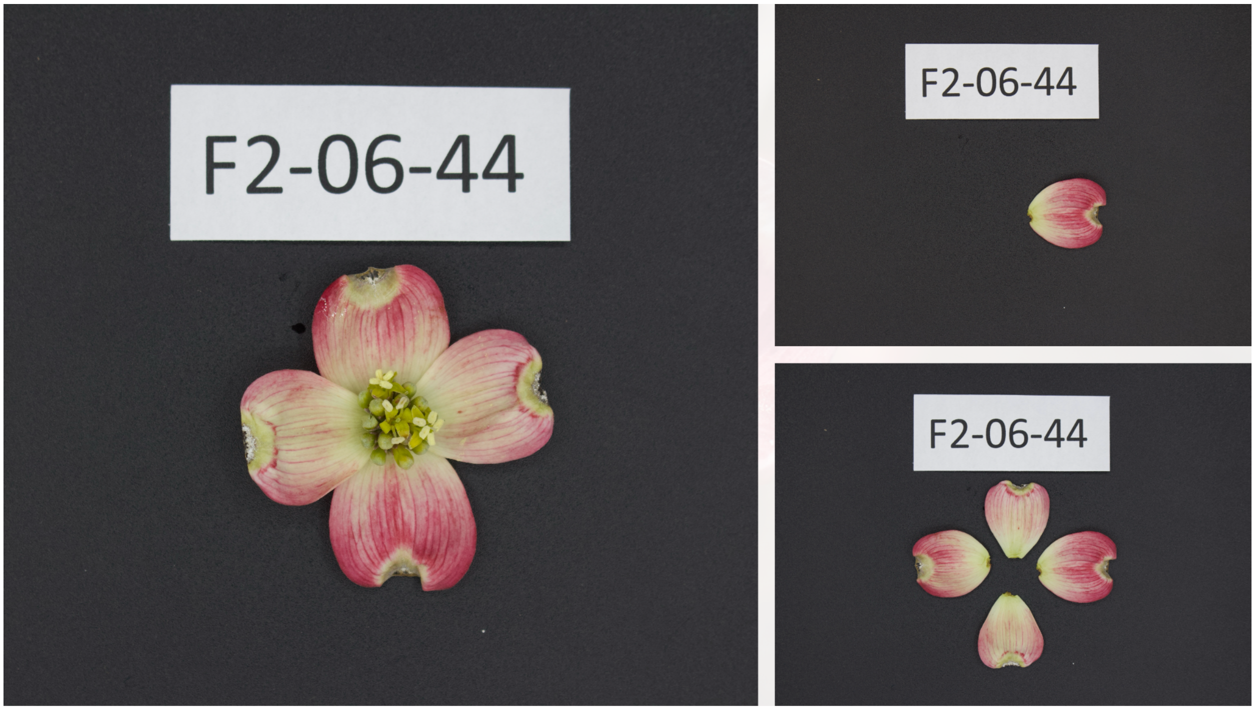


**Figure S1.** Examples of the three types of images captured for phenotyping the "pseudo-F2" population of flowering dogwoods. Left image: full inflorescence image, Right top: one bract image, Right bottom: four-bract images.

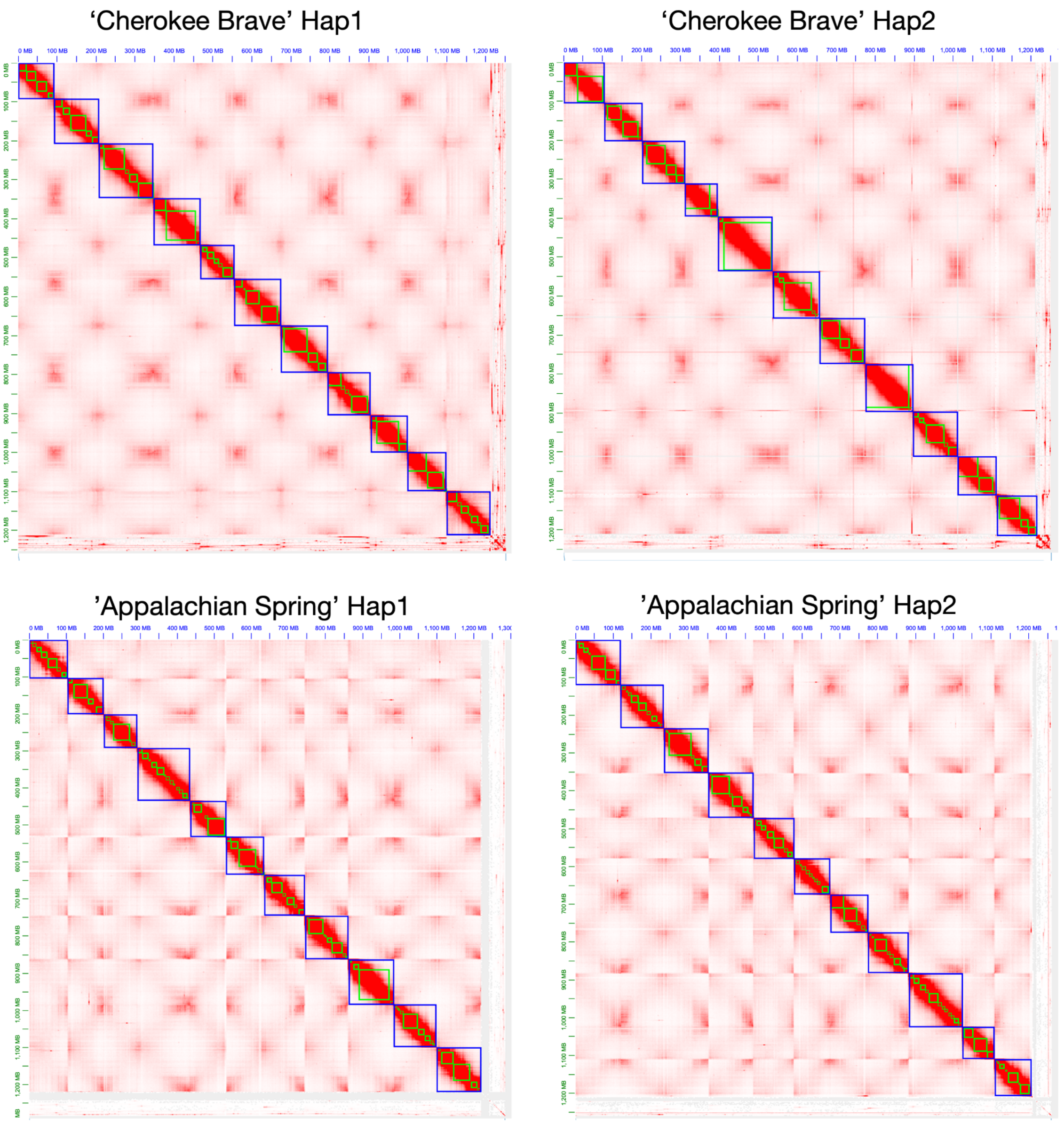


**Figure S2.** Hi-C contact maps for the four chromosome scale assemblies of flowering dogwood.

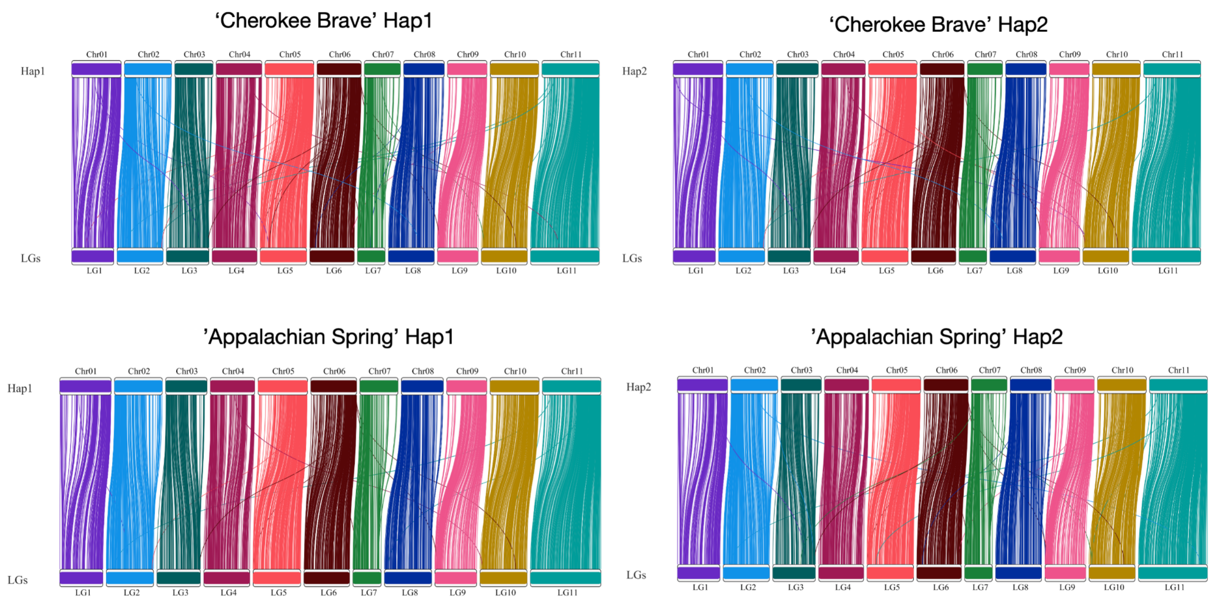


**Figure S3.** Visualization of the P25×P28 linkage map genetic markers from Moreau et al., 2022 mapped to the four flowering dogwood genome assemblies.

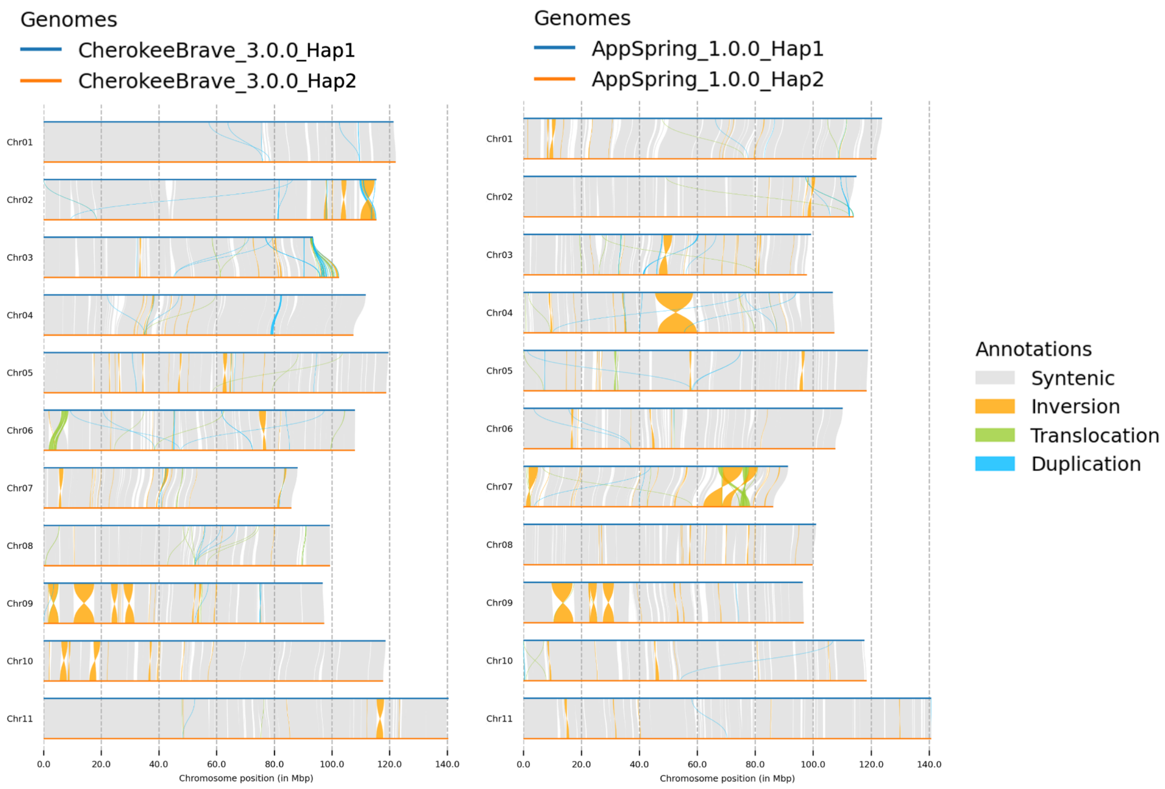


**Figure S4.** SyRI plot comparing haplotype 1 and 2 for *Cornus florida* 'Cherokee Brave' (left) and *C. florida* 'Appalachian Spring' (right) genomes.

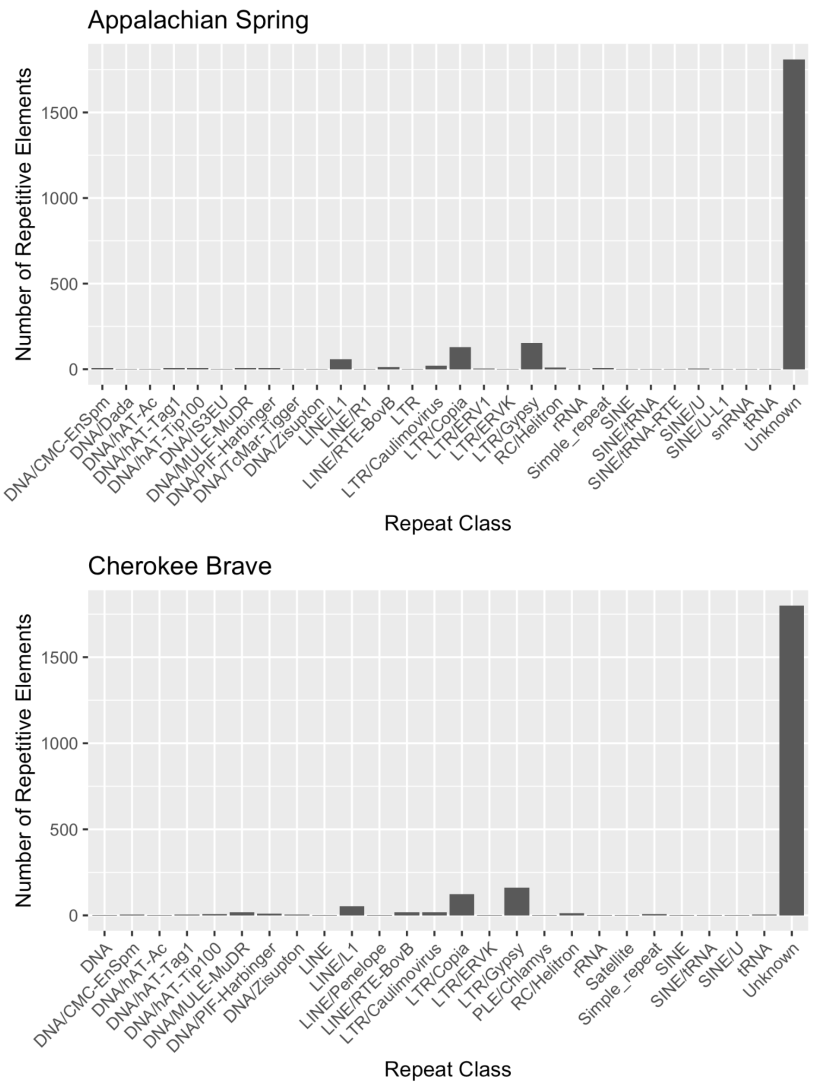


**Figure S5.** Repeat classes soft-masked in *Cornus florida* 'Appalachian Spring' and 'Cherokee Brave' genomes.

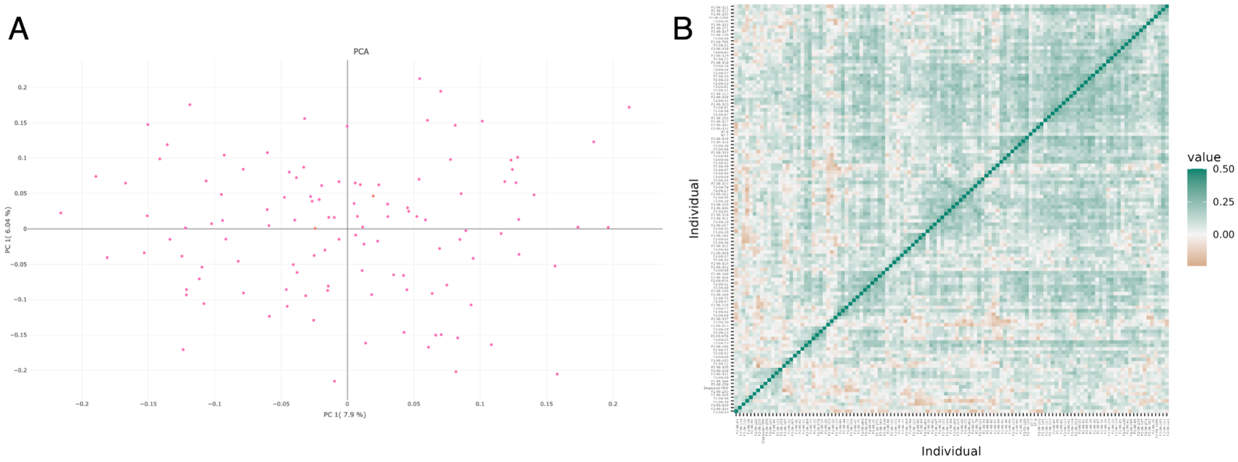


**Figure S6.** A) PCA of SNPs filtered to maximum missing of 20% and minor allele frequency of 0.05. B) Kinship matrix of all 125 individuals in *Cornus florida* 'Appalachian Spring' × 'Cherokee Brave' "pseudo-F2" plot. Brown squares indicate a negative kinship value (i.e., unrelated).

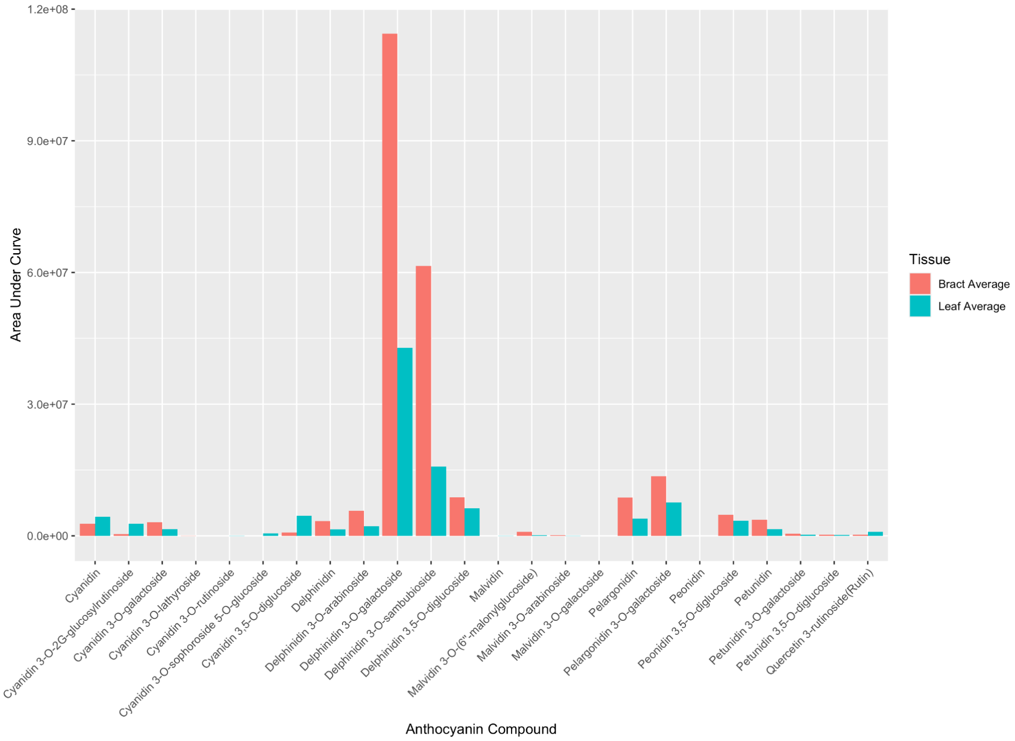


**Figure S7.** Anthocyanin profiling results of *Cornus florida* 'Cherokee Brave' leaves and bracts. Two samples for each tissue type were profiled and results were averaged.

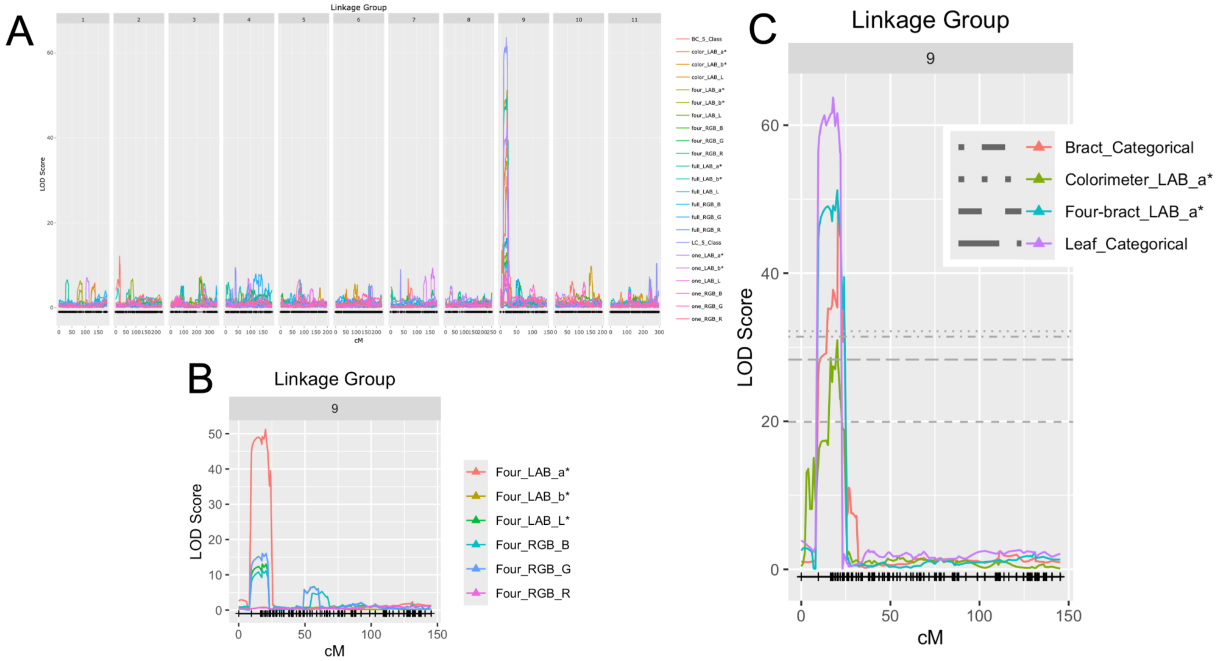


**Figure S8.** *Cornus florida* 'Appalachian Spring' x 'Cherokee Brave' "pseudo-F2" flowering dogwood QTL results of A) all 23 traits across the entire genome B) all four-bract image traits for LG9 C) colorimeter a*, four-bract image a*, bract categorical, and leaf categorical with respective permutation thresholds. Note that the population was treated as an F1 outcrossing population using 97-6 and 97-7 (the F1s of 'Appalachian Spring' and 'Cherokee Brave') as the parents.

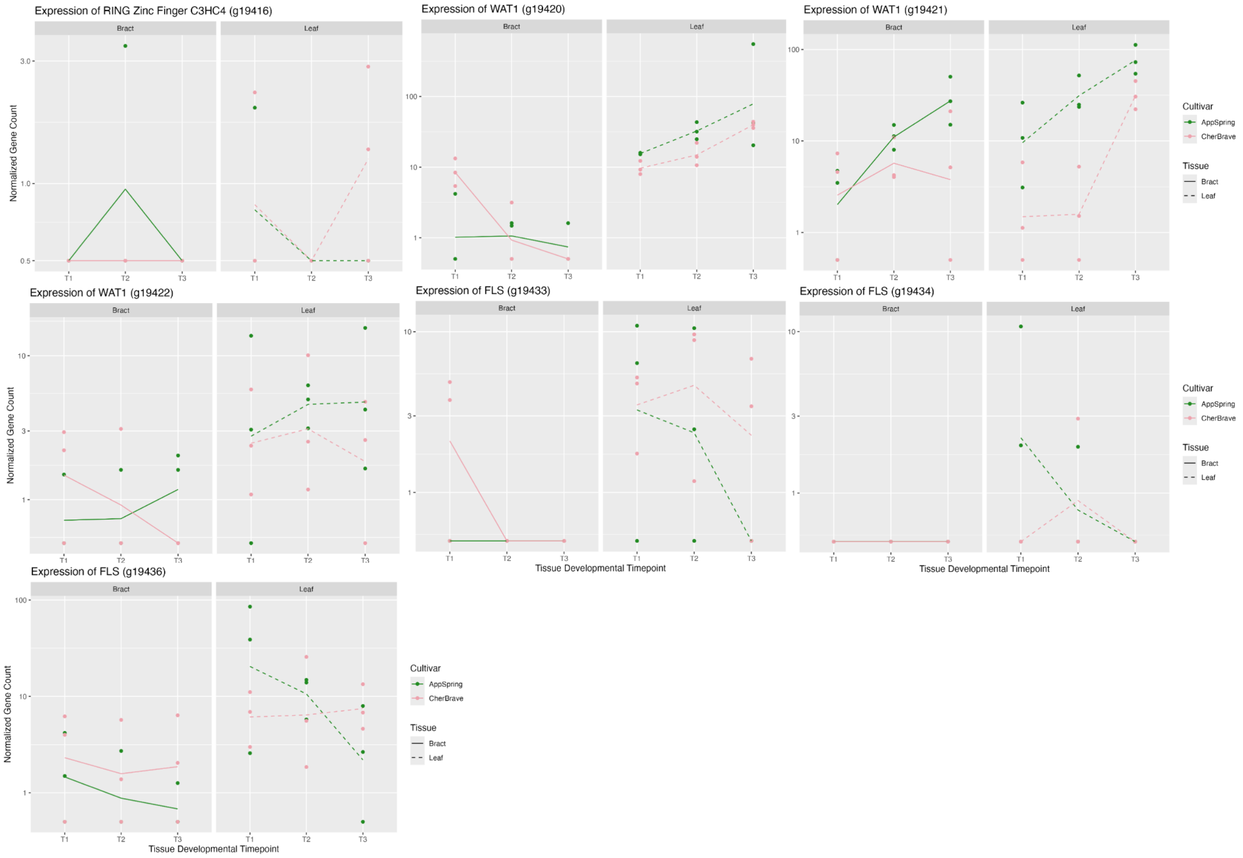


**Figure S9.** Expression of genes in 'Cherokee Brave' and 'Appalachian Spring' on a developmental time course. T1 = breaking bud, T2 = expanding tissue, T3 = fully expanded tissue. 'Appalachian Spring' has green leaves, white bracts, and gene counts are in green. 'Cherokee Brave' has red leaves, pink bracts, and gene counts are in pink. Gene counts in the bracts have a solid line and gene counts in the leaves have a dashed line.

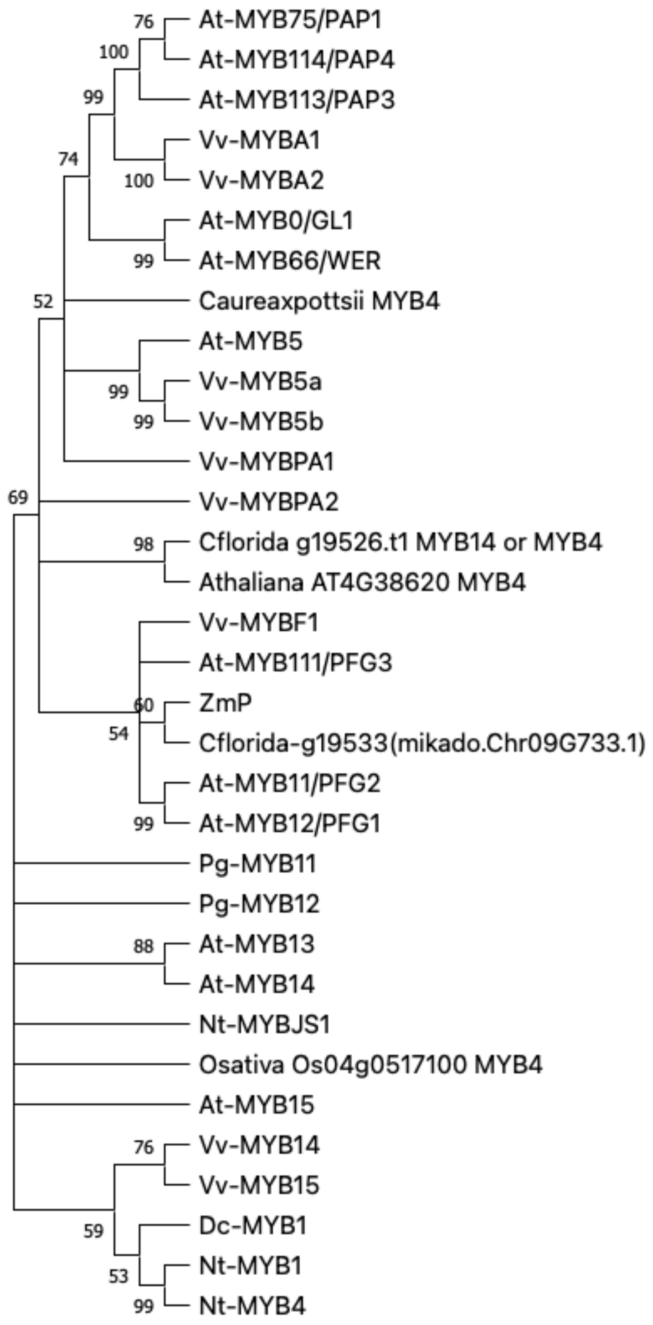


**Figure S10.** Neighbor-joining phylogenetic tree of MYB transcription factors. Bootstrap support is labeled. Gene of interest, g19533(mikado.Chr09G733.1) grouped with ZmP, a MYB-related protein in maize with bootstrap support of 60. Gene of interest, g19526, grouped with Arabidopsis MYB4 with bootstrap support of 98.

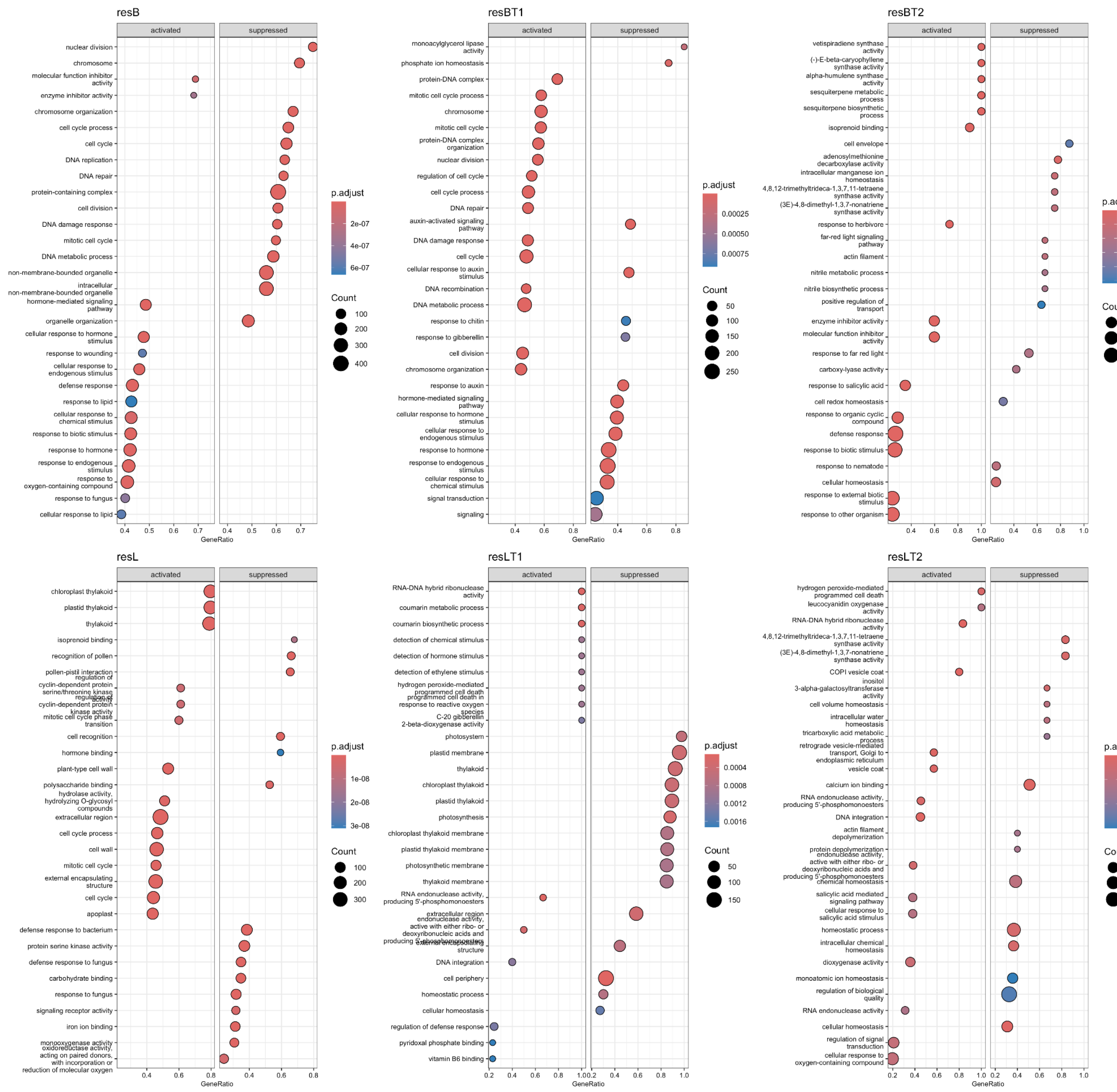


**Figure S11.** Activated and suppressed enriched GO terms in the overall bract and leaf time course RNAseq experiment (column 1), timepoint 1 (column 2), and timepoint 2 (column 3).

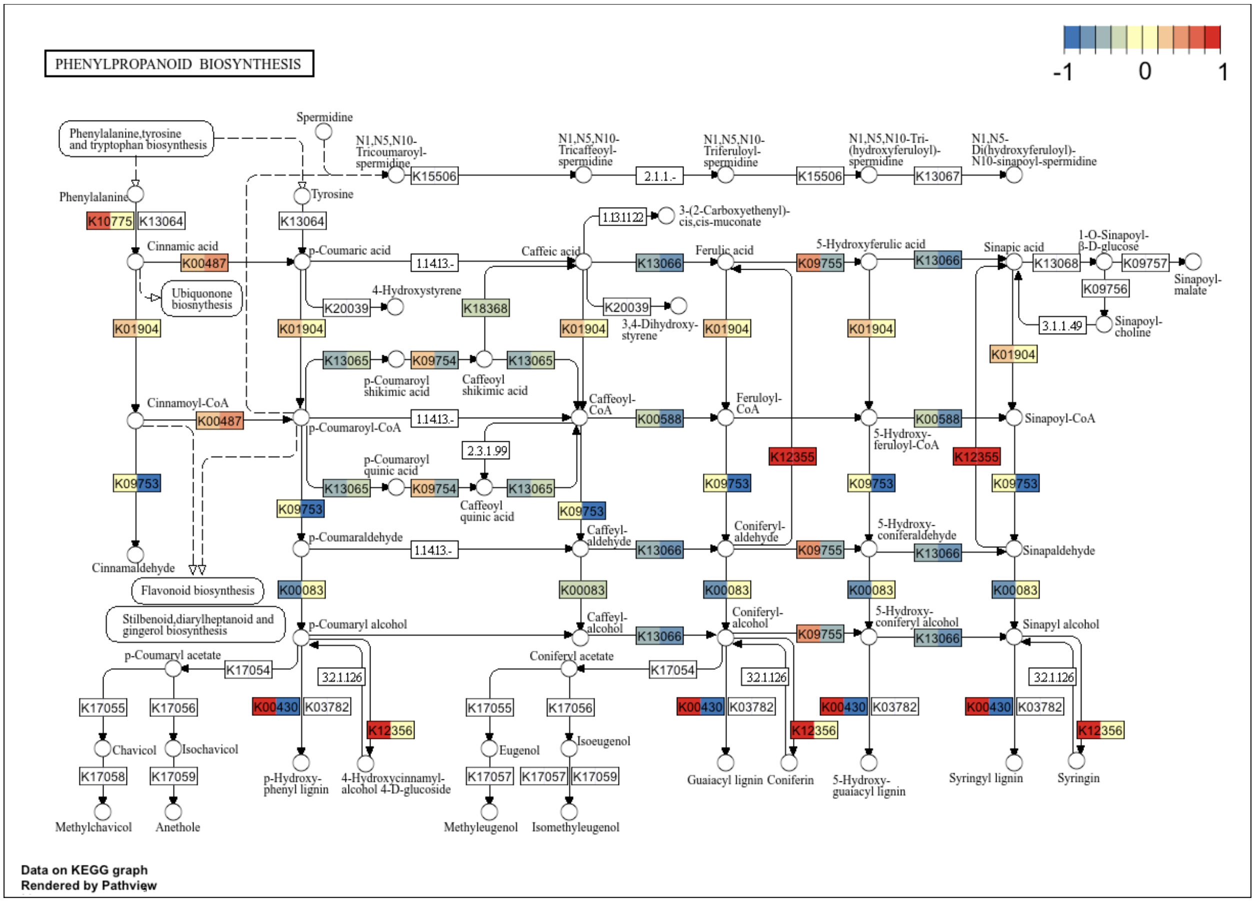


**Figure S12.** Maximum likelihood estimates of log2fold change of cultivar-specific gene expression within the KEGG phenylpropanoid biosynthetic pathway for timepoint 1 of time course experiment. Log2fold change in bracts is on the left and leaves are on the right.

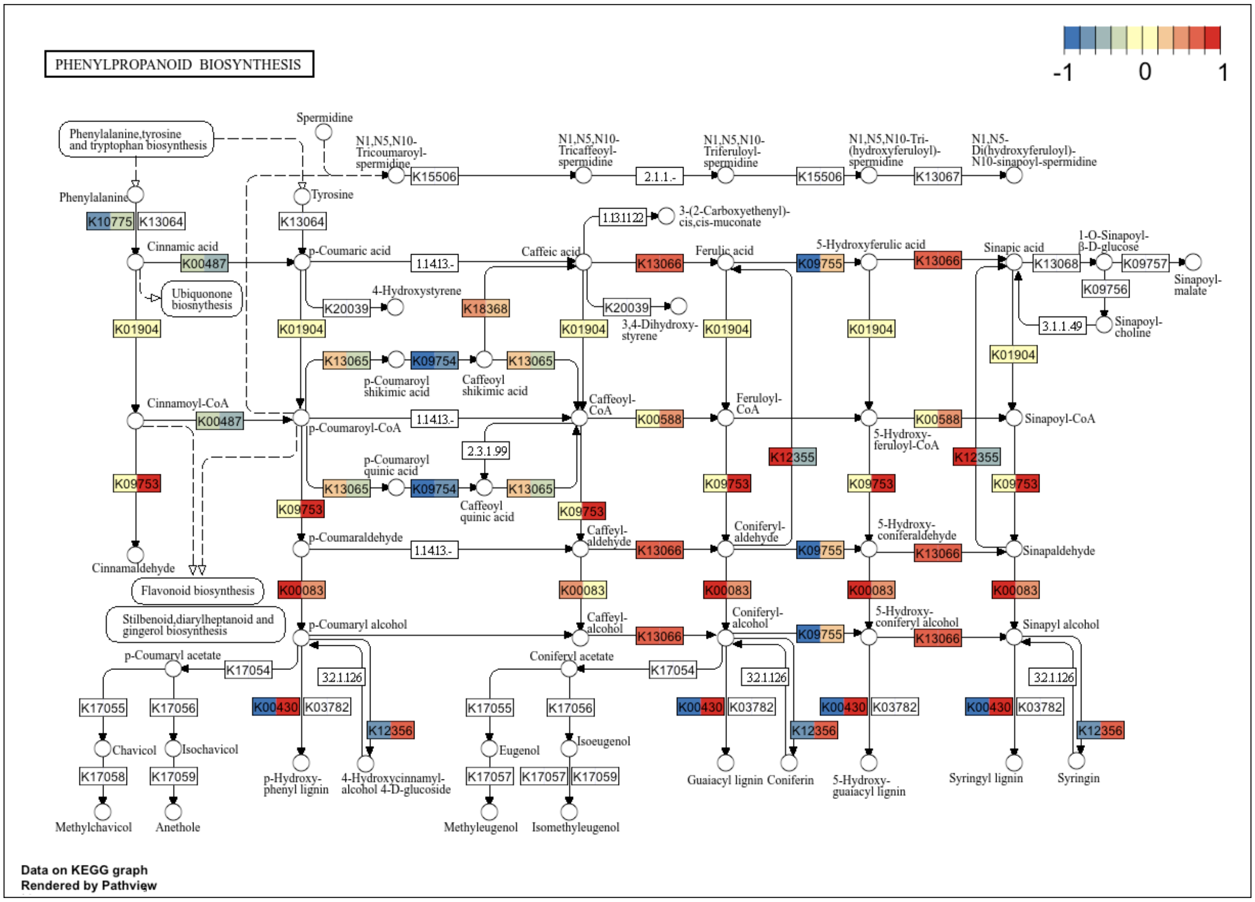


**Figure S13.** Maximum likelihood estimates of log2fold change of cultivar-specific gene expression within the KEGG phenylpropanoid biosynthetic pathway for timepoint 2 of time course experiment. Log2fold change in bracts is on the left and leaves are on the right.

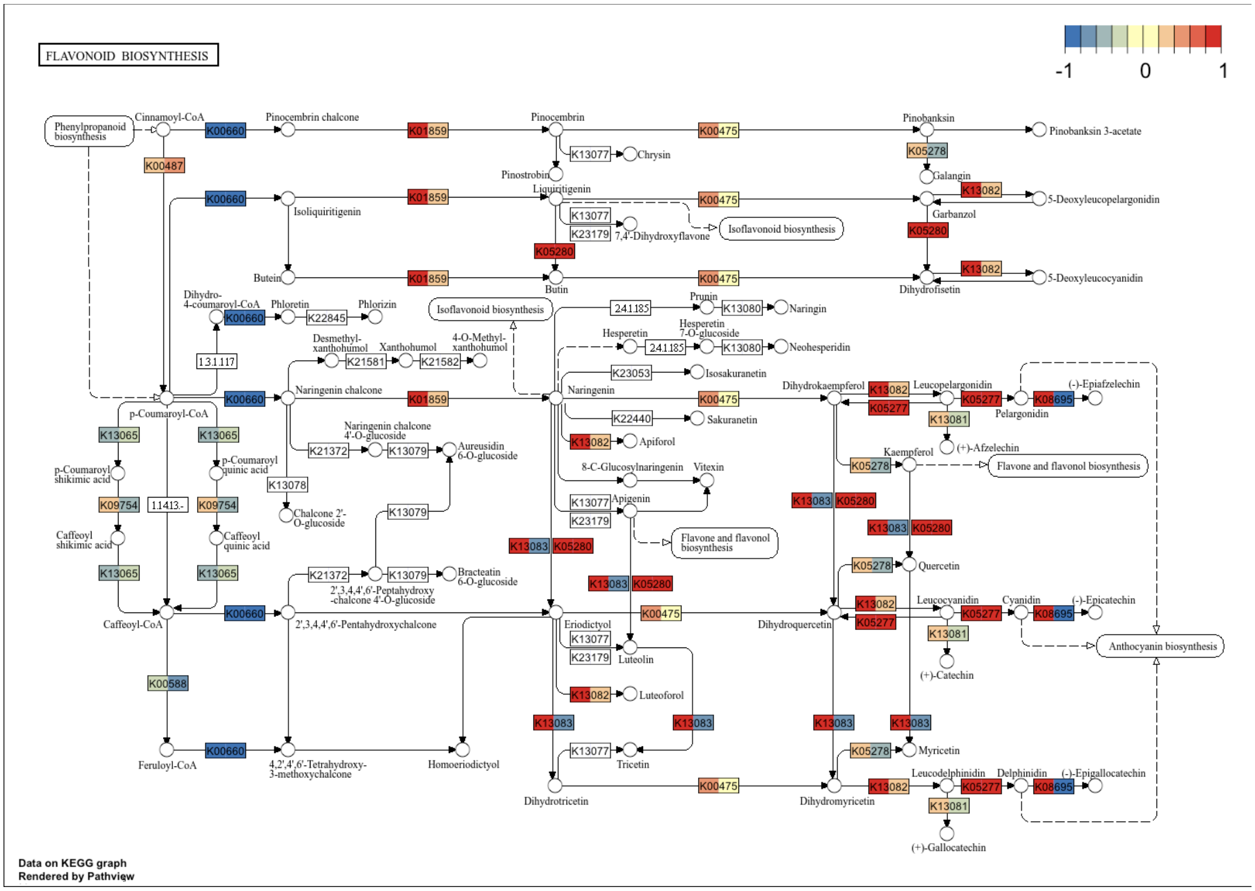


**Figure S14.** Maximum likelihood estimates of log2fold change of cultivar-specific gene expression within the KEGG flavonoid biosynthetic pathway for timepoint 1 of time course experiment. Log2fold change in bracts is on the left and leaves are on the right.

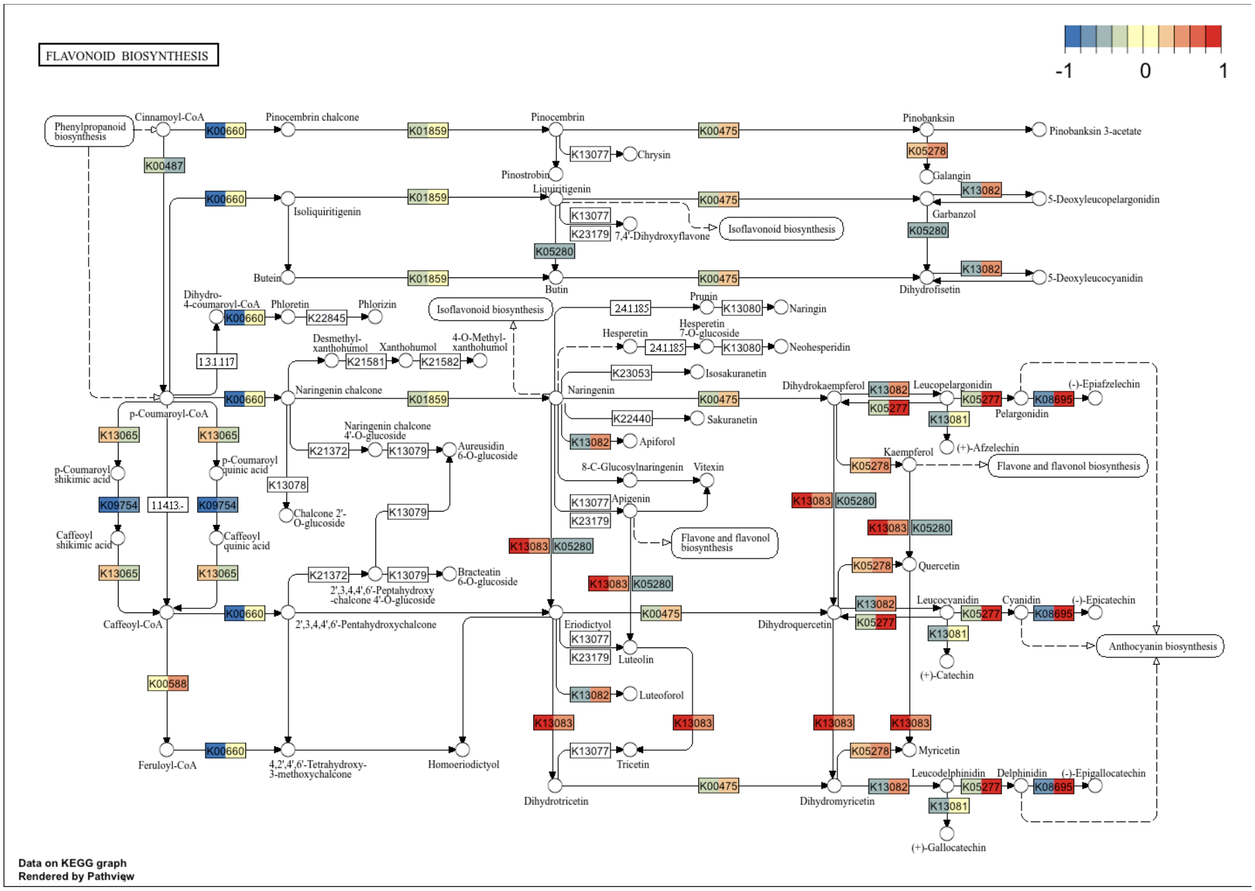


**Figure S15.** Maximum likelihood estimates of log2fold change of cultivar-specific gene expression within the KEGG flavonoid biosynthetic pathway for timepoint 2 of time course experiment. Log2fold change in bracts is on the left and leaves are on the right.

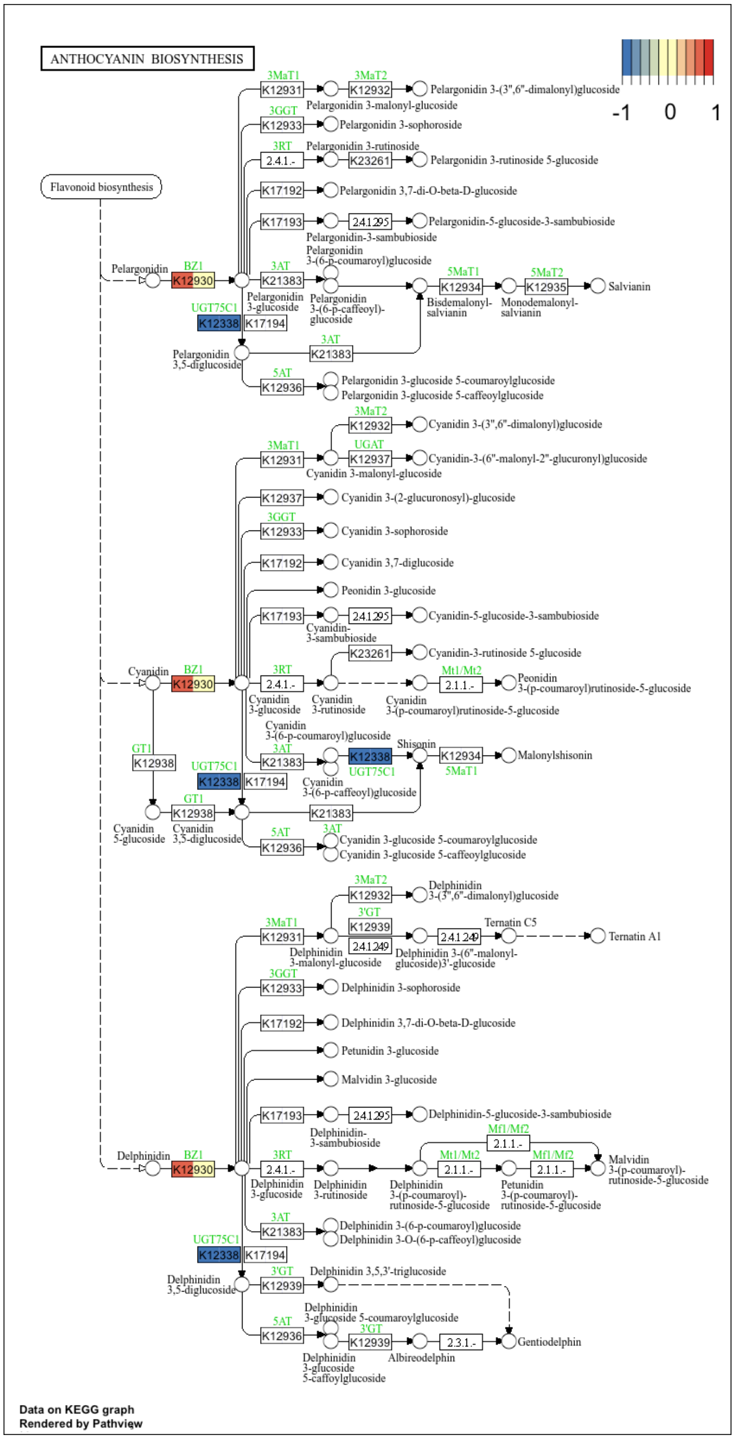


**Figure S16.** Maximum likelihood estimates of log2fold change of cultivar-specific gene expression within the KEGG anthocyanin biosynthetic pathway for timepoint 1 of time course experiment. Log2fold change in bracts is on the left and leaves are on the right.

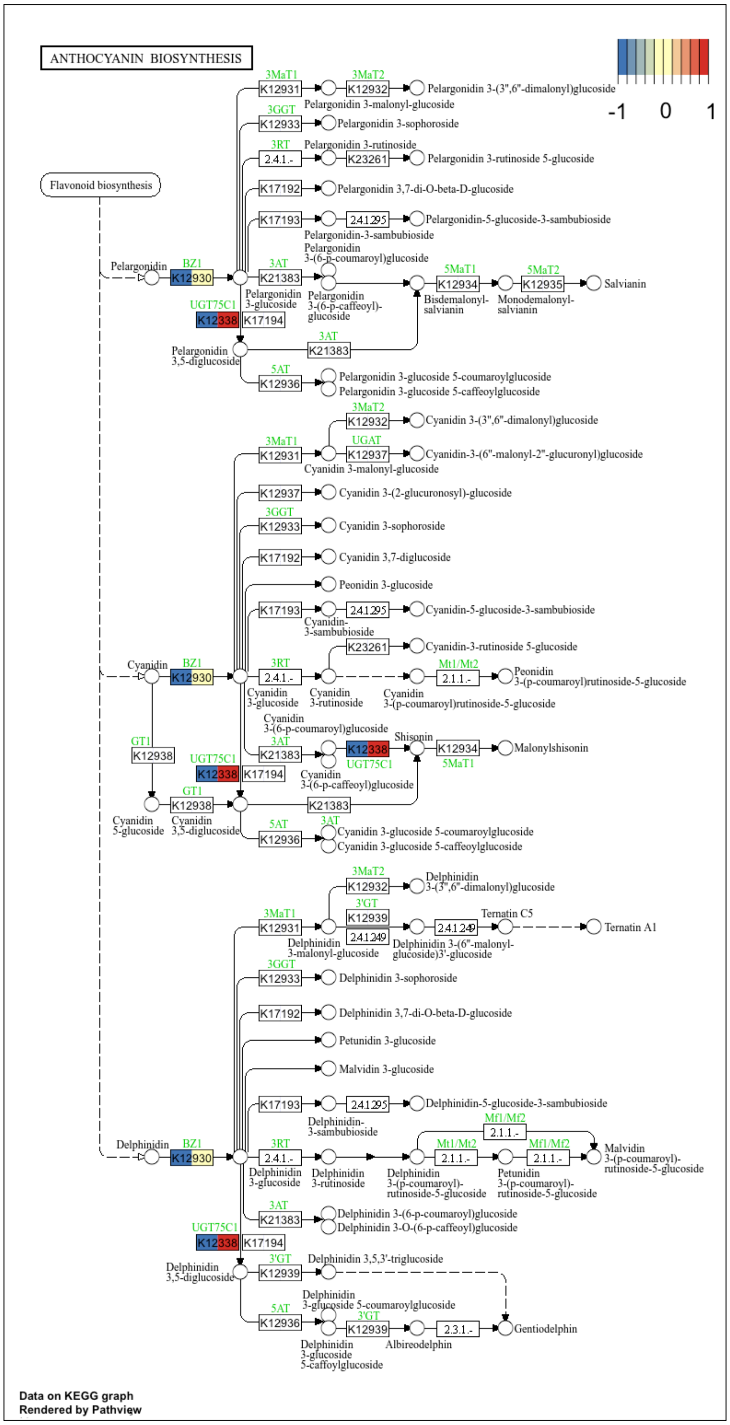

**Figure S17.** Maximum likelihood estimates of log2fold change of cultivar-specific gene expression within the KEGG anthocyanin biosynthetic pathway for timepoint 2 of time course experiment. Log2fold change in bracts is on the left and leaves are on the right.

**Methods S1** Supplementary Methods.

*Plant Materials and Sequencing–Genome Assembly*

Flowering dogwood cultivars 'Cherokee Brave' and 'Appalachian Spring' were selected for Pacific Biosciences HiFi and Illumina Hi-C sequencing and subsequent genome assembly. 'Cherokee Brave' flower buds were collected from a residential tree in Knoxville, TN, U.S. and frozen in a -80°C freezer. Bionano Prep Plant Tissue DNA Isolation kit (Bionano Genomics, San Diego, CA, U.S.) was used to isolate nuclei from the bud tissue. High molecular weight (HMW) DNA was extracted from the nuclei with Circulomics Nanobind Plant Nuclei Big DNA Kit (Pacific Biosciences, Menlo Park, CA, U.S.). HMW DNA was sheared to ~15kb fragments with the Covaris g-TUBE (Woburn, MA, U.S.). The sheared DNA was used to prepare a PacBio SMRTbell library and size selected using BluePippin (Sage Science, Beverly, MA, U.S.). The library was sequenced on three Sequel IIe (Pacific Biosciences, Menlo Park, CA, U.S.) cells. The PacBio HiFi reads are available through the NCBI sequence read archive (SRA) under BioProject PRJNA988529. Genome size was confirmed with flow cytometry. Nuclei were isolated from young 'Cherokee Brave' and pea (*Pisum sativum*) leaves using an LB01 + DAPI buffer. The amount of DNA within the nuclei was measured at the University of Georgia CTEGD Cytometry Shared Resource Laboratory. Young leaves from 'Appalachian Spring' were collected on the campus of the University of Georgia, and HMW DNA was extracted with the PacBio Circulomics Nanobind Kit. The library was sequenced on four Sequel II cells. The PacBio HiFi reads for 'Appalachian Spring' are available under BioProject PRJNA988529.

Young leaves were collected from 'Appalachian Spring' and 'Cherokee Brave' trees on the University of Tennessee, Knoxville campus and shipped wrapped in wet paper towels, on ice packs to Phase Genomics (Seattle, WA, U.S.) for Hi-C sequencing. The Hi-C reads are available under BioProject PRJNA988529.

*Plant Materials and Sequencing–Genotyping*

A "pseudo-F2" population was previously generated from 'Cherokee Brave' and 'Appalachian Spring' (Wang *et al.*, 2009). Due to the self-incompatibility of flowering dogwood and a second generation being necessary to recover the segregating pink bract phenotype, two ‘Cherokee Brave’ × ‘Appalachian Spring’ F1s (97-7 and 97-6) were crossed to generate the “pseudo-F2” population (Wang *et al.*, 2009). This population segregates for bract and leaf color and remains at the University of Tennessee Arboretum (Oak Ridge, TN, U.S.). To genotype the population, young leaf material was collected from all 125 individuals in the population. DNA was extracted using the QIAcube (Qiagen, Hilden, Germany) with the QIAamp 96 DNA QIAcube HT Kit (Qiagen) following the manufacturer's protocol, except for doubling the RNase (8ul) and adding 2% mass/volume polyvinylpyrrolidone (PVP) to the working solution. If the samples failed QC, they were tried again on the QIAcube. If the sample failed again, they were hand extracted with E.Z.N.A. Plant DNA DS Kit (Omega Biotek, Norcross, GA, U.S.). The manufacturer's protocol was followed, except 2% PVP was added to the first buffer. Samples were tested for quality by running gDNA on a 1% agarose gel, Qubit Flex Fluorometer (Thermo Fisher Scientific, Waltham, MA, U.S.), NanoDrop Lite Spectrophotometer (Thermo Fisher Scientific), and digestion with restriction enzymes. Once quality was ensured, samples were sent for genotype-by-sequencing (GBS) at The University of Wisconsin-Madison Biotechnology Center (Madison, WI, U.S.) with *PstI*/*MspI* restriction enzymes. With only 125 samples, only one and a half 96-well plates were used for sequencing. The individuals in the second partial plate had fewer reads than the first plate, therefore, a second round of sequencing was completed for the second partial plate. The parents (97-6 and 97-7) were sequenced multiple times to ensure greater sequencing depth. The demultiplexed GBS reads are available under BioProject PRJNA988529.

*Plant Materials and Sequencing–RNAseq*

To identify differentially expressed genes between the white and pink bracts as well as the green and red leaves, developing leaf and bract tissue were collected and immediately flash frozen in liquid nitrogen. Three replicates of tissues were collected from one ‘Cherokee Brave’ (located in a residential area in Knoxville, TN, U.S.) and two ‘Appalachian Spring’ (located on the University of Tennessee, Knoxville campus, U.S.) cultivars over a time course (Figure 1a), for a total of 36 samples. Tissue was collected at the following three developmental timepoints: 1) breaking bud, 2) expanding tissue, and 3) fully expanded tissue. Samples were ground and homogenized with a Beadmill 24 (Fisher Scientific, Pittsburgh, Pennsylvania, U.S.). If more than one 30 s cycle was needed, samples were put back in liquid nitrogen for 5 min before homogenization again. Total RNA was extracted from the bract and leaf tissues using a modified CTAB protocol (Doyle, 1990). Quality was checked using a 2100 Bioanalyzer (Agilent, Santa Clara, CA, U.S.) with the RNA 6000 Nano kit (Agilent).

Twenty-three of the RNAseq libraries were prepared by the UTIA Genomics Center for the Advancement of Agriculture (Knoxville, TN, U.S) using the SMART-Seq mRNA LP kit (Takara Bio, Kusatsu, Japan) and sequenced at the UT Genomics Core (Knoxville, TN, U.S.) using a NovaSeq 6000 (Illumina, San Diego, CA, U.S.). The other 13 samples were prepared and sequenced at Novogene (Sacramento, CA, U.S.) using their Illumina NovaSeq 6000 PE 150-bp.

*Genome Assembly and Scaffolding*

Chromosome-scale, partially haplotype-resolved genome assemblies were completed for both 'Cherokee Brave' and 'Appalachian Spring' using default parameters of hifiasm 0.16.1-r375 (Cheng *et al.*, 2021). QC checks were completed with stats.sh in BBTools (Bushnell, 2022) and BUSCO 5.3.2 using the embryophyta_odb10 dataset run in genome mode (Manni *et al.*, 2021). Both the HiFi and Hi-C data were used as input to build this initial contig-level assembly. From there, Hi-C contacts were identified in the preliminary hifiasm assembly with juicer 1.6 (Durand *et al.*, 2016b). *DpnI* restriction sites were used to generate site positions. The genome was then scaffolded into 11 chromosomes using 3D-DNA 201008 (Dudchenko *et al.*, 2017) with these parameters: `--editor-repeat-coverage 8 --editor-coarse-resolution 50000 --editor-coarse-region 250000`. QC checks were performed again using BBTools and BUSCO.

The genome was numbered and oriented against an already existing linkage map for flowering dogwood (Pfarr Moreau *et al.*, 2022). Loci were blasted against the genome using NCBI blast 2.11.0 (Camacho *et al.*, 2009), and to ensure correct orientation after flipping, blast hits were visualized with RIdeogram 0.2.2 (Hao *et al.*, 2020) on RStudio 2024.04.2+764 using R version 4.4.1. To subjectively assess the quality of the genomes, haplotypes 1 and 2 for each cultivar were compared to one another. This assessment was completed by aligning the two haplotypes with minimap2 (2.24-r1122) -ax asm5 –eqx (Li, 2021). Syntenic regions and structural variations were then identified using SyRI 1.7.0 (Goel *et al.*, 2019) and visualized with plotsr 1.1.1 (Goel & Schneeberger, 2022).

The scaffolded assemblies were edited with Juicebox 1.7.0 (Durand *et al.*, 2016a) by inverting contigs that appeared to be inversions in both the Hi-C contact map and in the plotsr output, as well as moving contigs to debris if there were limited Hi-C contacts to support the contig placement. The resulting assembly file was then used with run-asm-pipeline-post-review.sh script in 3D-DNA to make the changes to the fasta files. SyRI was used again to confirm that large inversions were fixed.

Each haplotype was then aligned against the flowering dogwood chloroplast genome (NCBI: NC_044820.1) and rhododendron mitochondrial genome (NCBI: OQ450179.1), and any scaffolds with >90% coverage were removed from the assembly. The NCBI Foreign Contamination Screens, including FCS-gx 0.4.0 and FCS-adaptor 0.5.0 (Astashyn *et al.*, 2024), were then run and any scaffolds identified as contamination were removed. One final round of QC was then completed with BBTools and BUSCO.

*Genome Annotation*

Repeats were identified with RepeatModeler 2.0.3 (Flynn *et al.*, 2020) and masked using RepeatMasker 4.1.3-p1 (Smit *et al.*, 2013) with the Eudicot repeats from Dfam.h5 (Hubley *et al.*, 2016), Repbase 20181026 (Bao *et al.*, 2015), and output from RepeatModeler. For repeat identification with RepeatModeler and repeat masking with RepeatMasker, the haploid assemblies for each individual were combined (i.e., repeat identification and masking were completed on the diploid assemblies). After masking, the genomes were split back into separate haploid files. Raw RNAseq reads from 5 SRA leaf samples (‘Appalachian Spring’: SRR10616261, SRR10616272, SRR10616244, SRR10616275, and SRR10616251; ‘Cherokee Brave’: SRR10584713, SRR21411472, SRR21411471, SRR10584708, and SRR10584701) and 3 bract and 2 leaf samples (‘Appalachian Spring’ and ‘Cherokee Brave’ South samples; sequencing details above) for each cultivar were aligned to the soft-masked genomes using Star 2.7.6a (Dobin *et al.*, 2013). Other extrinsic evidence that was fed into BRAKER3 (Gabriel *et al.*, 2023) was the Viridiplantae orthoDB v11 protein database (Kuznetsov *et al.*, 2023) with the proteomes of *Hydrangea quercifolia* v1.0 (Cornales; 589), *Vaccinium darrowii* v1.2 (Ericales; 700), *Lindenbergia philippensis* v1.1 (Lamiales; 689), *Mimulus guttatus* TOL v5.0 (Lamiales; 551) from JGI Phytozome 13 (Goodstein *et al.*, 2012). Stop codons indicated with '*' were removed from the dataset. The BRAKER3 3.0.3 (docker container v.1.0.3) was used with the –gff3 flag to annotate all genomes.

Before evaluating annotation completeness, only the longest transcript was retained in the fasta file. Using the single-transcript fasta file, BUSCO in protein mode was used to evaluate how complete the annotations were, and EnTAP 1.0.0 (Hart *et al.*, 2020) was used to functionally annotate the genes and evaluate the proportion of functionally annotated genes. gFACs was used in `–false-run' mode to calculate the mono:multi exonic ratio for the annotated genes (Caballero & Wegrzyn, 2019).

*Population Genotyping*

To genotype the 125 individuals in the "pseudo-F2" population, the raw GBS reads were demultiplexed with sabre 1.0 (Joshi, 2011). Fastp 0.23.4 was used for adapter removal and trimming the demultiplexed reads (Chen, 2023). The trimmed reads were aligned to the 'Cherokee Brave' Haplotype 2 genome (this was used as the reference throughout the rest of this study) with bwa 0.7.17 (Li, 2013) and sorted with samtools v.1.15.1 (Danecek *et al.*, 2021). Joint genotyping was completed with GATK 4.2.2.0 (Poplin *et al.*, 2017), and the resulting SNPs were flagged using `gatk VariantFiltration`, following GATK guidelines (QD < 2.0, FS > 60.0, MQ < 40.0, SOR > 4.0, MQRankSum < -12.5, and ReadPosRankSum < -8.0). Failed SNPs were filtered out, and one sample that failed sequencing was removed from the dataset at this point using bcftools 1.6 (Danecek *et al.*, 2021). The dataset was filtered for a minimum read depth of 10 and varying maximum missing levels (5-20%) and minor allele frequency (0.05-0.2). The resulting VCF files were visualized with principal component analysis (PCA), calculated with SNPRelate 1.40 (Zheng *et al.*, 2012), to ensure no bias could be detected between the two plates sequenced. A kinship matrix was built using `--make-king 'square'` flag of plink2 2.00 (Chang *et al.*, 2015) and visualized with a heatmap in R with ggplot 3.5.1. Eight samples that had an average kinship value of <0 were removed from the dataset (F2-06-01, F2-06-120, F2-06-

220A, F2-06-128, F2-06-63, F2-06-82, F2-06-60, and F2-06-219). The VCF file filtered for a maximum missing rate of 20% and minor allele frequency of 0.05 was used for linkage map construction.

*Linkage Map Construction*

Linkage map construction was completed with OneMap 3.0.0 (Margarido *et al.*, 2007) using both the recombination and reference genome information of the individuals left after SNP and relatedness filtering (n=116). The tutorial vignette for outcrossing populations was followed, using ‘97-6’ as parent 1 and ‘97-7’ as parent 2. Despite this population being a “pseudo-F2” population of ‘Cherokee Brave’ × ‘Appalachian Spring,’ it was treated as a F1 population of ‘97-6’ × ‘97-7’ for linkage map construction and QTL analysis due to the heterozygosity of flowering dogwood. The function map_avoid_unlinked was used for original map estimation. The recombination matrix heatmaps for each linkage group were then manually inspected to remove markers that did not have the expected color pattern (i.e., <0.1 recombination fractions/red along the diagonal). The map order was estimated again after marker removal. Multiple map order functions were tested for each linkage group. The function map_avoid_unlinked produced maps with the smallest map size (cM) and the largest log-likelihood, except LG03, in which the ug function was used. Each marker was manually added back to the map to see if its position in the map changed or improved the log-likelihood of the map.

*Population Phenotyping*

The “pseudo-F2” population was phenotyped between March 31 and April 6, 2023, at the UT Arboretum (Oak Ridge, TN, U.S.). Due to bract color being indicative of leaf color and bract color being the more important phenotypic trait, characterizing bract color was prioritized over leaf color for this experiment. Therefore, only categorical data was collected for the leaves. Trees (n=125) were surveyed every day during this period, and samples were collected when most inflorescences on a tree had only one flower open. Eight trees did not bloom that year, and therefore, the only phenotypic information collected was the categorical leaf color. For collection (n=117), the three inflorescences with maximal color and least amount of spot anthracnose were placed into a plastic bag with a wet paper towel and a tree label. Samples were placed in a cooler with ice packs until all samples were collected for the day. Multiple phenotyping methods were used. The inflorescences were photographed in a FotodioX LED Studio-In-a-Box (Gurnee, IL, U.S.) with a Canon EOS Rebel T5i camera with a Canon EF-S 18-55mm f/3.5-5.6 IS STM lens (Ota City, Tokyo, Japan). Images were taken in manual mode with a focal length of 55mm, f-number of f/5.6, an exposure time of 1/250, and saved in RAW format. For each inflorescence collected, three images were captured (Figure S1.): 1) full inflorescence, 2) four bracts pulled off of true flowers, and 3) one bract. The one bract in image 3 also had two colorimeter readings taken with the CR-10 Plus Colorimeter (Konica Minolta, Chiyoda, Tokyo, Japan). Measurements were taken in the L*a*b* color scale (L = lightness, a* = red-green axis, and b* = yellow-blue axis) on each side of the bract. Categorical data was also collected, for leaves and bracts, using five classes (1= white bracted/green leafed, 2 = less than half pink/red, 3 = half pink/red, 4 = more than half pink/red, 5 = almost completely pink/red; Figure 1).

The white balance of the images was corrected using RawTherapee (https://rawtherapee.com/), Color > White Balance > Method: Custom > Pick, and using the dropper, the white label in the photo was selected. To treat every image the same, the .pp3 file that was created doing this for the first image was applied to all images. Images were exported as .tiff. A custom Python script was used to rename images with the sample name (available https://github.com/trinityhamm/Flowering-Dogwood-Color). Images were read in using Pillow 9.4.0 (Clark, 2023). The Python distribution of Tesseract OCR (pytesseract 0.3.10) (Hoffstaetter, 2024) was used with the image_to_string function to pull tree IDs out of the images. This was not always successful, so multiple configurations were iteratively used to maximize the label removal. Within Pillow, the images were either read in as is or converted to grey scale using the 'LA' or 'L' method. Within pytesseract, page segmentation options (–psm) 1) Automatic page segmentation with OSD and 12) Sparse text with OSD were used. The os.rename function was then used to rename the images with the output from pytesseract.image_to_string, as long as it started with 'F2'. Images were manually cross-checked, and any incorrect names were fixed. Images were then analyzed using a Fiji macro (Strock, 2021) to segment images to just the inflorescences or bracts and extract average RGB and L*a*b values. The only modification to the code was filter[1], which was modified to min[2]=50; max[2]=200; to aid in segmenting the image. The macro was run on the Full Inflorescence, Four Bract, and One Bract sets of images separately. The resulting .csv file from each was then processed with Python to average the RGB and L*a*b* values for each individual. The L*a*b* colorimeter readings were also averaged for each individual.

*Anthocyanin Profiling*

Anthocyanin profiling was performed on the bracts and leaves of 'Cherokee Brave'. Bracts and leaves with pink and red pigmentation were flash frozen and shipped to Creative Proteomics (Shirley, NY, U.S.) for qualitative anthocyanin profiling. Two separate leaf and bract samples were analyzed. The resulting area under the curve results were averaged for each tissue type.

*QTL Mapping*

All 23 phenotyping metrics (RGB and L*a*b* from full inflorescence, four bracts, and one bract images, L*a*b* from colorimeter, and five-class classification for the leaves and bracts) were used for QTL identification with fullsibQTL 0.0.9012 (Gazaffi *et al.*, 2020). A total of 116 individuals had leaf and genotypic data, and 108 individuals had bract, leaf, and genotypic data. Composite interval mapping (CIM) was performed with the cim_scan function. Cofactor selection was completed with cof_selection, which selected the markers to be used in the CIM scan based on Akaike Information Criterion (AIC). This was iteratively done for each phenotyping method. The threshold for QTL detection was then determined for select methods using the cim_scan with 1,000 permutations.

*Differential Gene Expression*

Identifying differentially expressed genes between the white-bracted/green-leafed 'Appalachian Spring' and the pink-bracted/red-leafed 'Cherokee Brave' began with assessing the quality of reads with fastqc 0.12.1 (Andrews, 2010). Raw reads were mapped to the soft-masked 'Cherokee Brave' Hap 2 genome with STAR 2.7.11b (Dobin *et al.*, 2013), using gene info from the BRAKER3 genome annotation. The aligned reads were then used to count transcripts with HTseq 2.0.4 (Anders *et al.*, 2015). Results were then read into DESeq2 1.46.0 (Love *et al.*,

2014) to identify differentially expressed genes. The experimental design included cultivar, sequencing location, timepoint, and the interaction between cultivar and timepoint. Data from the bracts and leaves were analyzed separately. 'Appalachian Spring' was the reference sample for both. To visualize, the original DESeq data set object was transformed with a regularized log, and a PCA was plotted. The likelihood ratio test (LRT) was used with a reduced model by removing the interaction term to identify genes that showed a cultivar-specific effect after time 0 (i.e., genes that move up or down the same way over time will not have small *p* values). Differentially expressed genes were identified for the overall experiment and specifically at timepoint 1 with the results function, using the Wald test at $\alpha = 0.05$.

An organism package for annotations was created for flowering dogwood using AnnotationForge 1.48 (Marc Carlson, 2025) and the EnTAP results. This database was used for gene enrichment analysis with clusterProfiler version 4.14.4 (Wu *et al.*, 2021). The proteome was annotated with KEGG Automatic Annotation Server (KAAS) (Moriya *et al.*, 2007). ClusterProfiler was also used for gene enrichment analysis with KEGG terms. To elucidate the identity of the discovered MYB TFs, a phylogenetic tree was constructed with the neighbor-joining method. Multiple sequence alignments of selected MYBs from GenBank or EMBL databases (Table S3.) were generated using clustalo (Sievers & Higgins, 2018) with 1000 iterations. Alignments were trimmed with trimAl (Capella-Gutiérrez *et al.*, 2009). The unrooted neighbor-joining tree was constructed with MEGA version 11 (Tamura *et al.*, 2021). During this analysis, the protein sequence of an identified MYB of interest (g19533) was compared to other plant MYB TFs and found to be truncated in length due to a gene annotation error. Using an in-house annotation pipeline, Ragnorak (https://github.com/ryandkuster/ragnarok), Chr09 was annotated, and the full protein sequence was recovered. The gene annotation from Ragnorak was used to construct the final phylogenetic tree.

*Contextualizing Results with Genomic Resources*

The four chromosome-scale assemblies (Cherokee Brave Hap 1 and 2–pink-bracted and red-leafed, and Appalachian Spring Hap 1 and 2–white-bracted and green-leafed) and their gene (from BRAKER3) and repeat (from RepeatModeler) annotations were visualized with

Persephone (Persephone Software, LLC). To better understand the locus of interest and validate differences, the HiFi reads for Cherokee Brave and Appalachian Spring were mapped back to the Cherokee Brave Hap2 genome, and the BAM file was visualized in tandem within Persephone. Additionally, the other three genomes were mapped back to the Cherokee Brave Hap2 genome with minimap2, and the PAF file was also visualized in tandem in Persephone. The VCF file was visualized with IGV 2.13, and SNPs with diagnostic potential were noted and pulled from the VCF with a Python script to calculate the frequency of predicting the phenotype correctly (available https://github.com/trinityhamm/Flowering-Dogwood-Color). The regions flanking the group of perfectly accurate diagnostic SNPs were inspected in Persephone, noting candidate genes relating to anthocyanin biosynthesis. For genes of interest, either from the list of differentially expressed genes, enrichment analysis, or visual inspection of the region and diagnostic SNPs, the normalized gene count throughout the experiment was calculated with the plotCounts() function with DESeq2 and visualized with ggplot.

**Persephone Software, LLC**. Persephone: A multi-genome browser carefully crafted using latest technologies.

**Pfarr Moreau E, Honig JA, Molnar TJ**. **2022**. High-density linkage mapping and identification of quantitative trait loci associated with powdery mildew resistance in flowering dogwood (*Cornus florida*). *Horticulturae* **8**: 405.

**Poplin R, Ruano-Rubio V, DePristo MA, Fennell TJ, Carneiro MO, Van der Auwera GA, Kling DE, Gauthier LD, Levy-Moonshine A, Roazen D**. **2017**. Scaling accurate genetic variant discovery to tens of thousands of samples. *BioRxiv*: 201178.

**Sievers F, Higgins DG**. **2018**. Clustal Omega for making accurate alignments of many protein sequences. *Protein Science* **27**: 135–145.

**Smit A, Hubley R, Green P**. **2013**. RepeatMasker Open.

**Strock C**. **2021**. Protocol for extracting basic color metrics from Images in ImageJ/Fiji.

**Tamura K, Stecher G, Kumar S**. **2021**. MEGA11: Molecular Evolutionary Genetics Analysis Version 11. *Molecular Biology and Evolution* **38**: 3022–3027.

**Wang X, Wadl PA, Rinehart TA, Scheffler BE, Windham MT, Spiers JM, Johnson DH, Trigiano RN**. **2009**. A linkage map for flowering dogwood (*Cornus florida* L.) based on microsatellite markers. *Euphytica* **165**: 165–175.

**Wu T, Hu E, Xu S, Chen M, Guo P, Dai Z, Feng T, Zhou L, Tang W, Zhan L, *et al.*2021**. clusterProfiler 4.0: A universal enrichment tool for interpreting omics data. *The Innovation* **2**.

**Zheng X, Levine D, Shen J, Gogarten SM, Laurie C, Weir BS**. **2012**. A high-performance computing toolset for relatedness and principal component analysis of SNP data. *Bioinformatics* **28**: 3326–3328.
